# Establishing Comprehensive Transthoracic Echocardiography Reference Ranges for Mouse Models: Insights into the Impact of Anesthesia, Sex, and Age

**DOI:** 10.1101/2025.04.14.647528

**Authors:** Manuela Oestereicher, Christopher Ward, Elida Schneltzer, Susan Marschall, Helmut Fuchs, Valerie Gailus-Durner, Ghina Bou About, Mohammed Selloum, Hamid Meziane, Michelle Stewart, Lydia Elizabeth Teboul, Clare Norris, Dale Pimm, Marina Kan, Federico Lopez Gomez, Robert Wilson, Mayra Monroy, Sheraz Pasha, Eva Zabrodska, Jan Prochazka, David Pajuelo Reguera, Zuzana Nichtova, Yann Herault, Sara Wells, Helen Parkinson, Jason Heaney, Radislav Sedlacek, Xiang Gao, Martin Hrabe de Angelis, Nadine Spielmann

## Abstract

Mouse models play a critical role in cardiology research, offering valuable insights into the molecular mechanisms, genetics, and potential treatments for cardiovascular diseases. However, the ability to transfer findings in mice between studies is limited by the absence of standardized protocols and valid reference values for the assessment of normal cardiac function in mice. This study aims to establish comprehensive transthoracic echocardiography (TTE) reference ranges for mice, particularly focusing on C57BL/6N wildtype controls. The study, which includes data from over 15,000 mice through the International Mouse Phenotyping Consortium (IMPC), highlights how variables such as sex, age, body weight, and anesthesia affect TTE parameters.

The findings showed that anesthesia is the primary predictor of variability in cardiac function. Isoflurane and tribromoethanol anesthetized mice presented with modified cardiac function compared to conscious mice. Additionally, we observed minimal sex differences in cardiac morphology and function, except for small variations influenced by anesthesia. The effects of aging on cardiac function were modest, characterized by a decrease in heart rate and subtle changes in ventricular dimensions without evidence of pathological remodeling, likely attributable to disease-free cardiovascular aging. Validation of the reference ranges across multiple mouse strains showed that these values provide a reliable baseline for experiments involving cardiac function in mice. The data underscore the importance of using anesthesia-specific reference values when interpreting TTE results, ensuring robust comparisons in genetic and pharmacological studies.

These reference ranges serve as quality assurance tools for future cardiac studies in mice, offering insights into typical TTE parameter values, supporting the detection of experimental perturbations, and contributing to more effective translation of findings from mouse to human.

## Introduction

Transthoracic echocardiography (TTE) is a vital non-invasive tool for diagnosing, monitoring, and guiding treatment in a wide range of heart conditions. TTE reference ranges from large healthy populations have become an increasingly common and valuable tool for diagnostic decision-making in clinical medicine (Lancellotti, Badano et al. 2013, Pfaffenberger, Bartko et al. 2013, Eriksen-Volnes, Grue et al. 2023, Nyberg, Jakobsen et al. 2023, Ahn, Koo et al. 2024).

Mouse models play a critical role in cardiology research, offering valuable insights into the molecular mechanisms, genetics, and potential treatments for cardiovascular diseases (Dobrev and Wehrens 2018, Lindsey, Brunt et al. 2021, Withaar, Lam et al. 2021). Their widespread use in cardiology stems from the ease of genetic manipulation in mice, their relatively short generation times and lifespans, and the similarity of many physiological and pathological processes between mice and humans (Zacchigna, Paldino et al. 2021).

Advancements in echocardiographic equipment and transducers have enabled its successful adaptation from humans to rodents (Gao, Ho et al. 2011, O’Riordan, Trochet et al. 2023). Hence, the establishment of TTE reference ranges healthy mice create a benchmark for normal cardiac physiology and will improve the understanding of normal values and variation across sex, age or strains.

TTE reference ranges are not only fundamental to evaluate disease models by revealing how diseases or interventions impact the model, but they also benchmark animal welfare by signaling health issues when values deviate – which can aid in assessment of drug safety, revealing potential toxic or adverse effects from normal values.

TTE reference ranges allow comparison across studies by standardizing data, ensuring consistency and reproducibility of research findings. In mouse models, they have the potential to improve data interpretation, animal welfare, and research reliability towards translatability (Robinson, Krieger et al. 2019, Domínguez-Oliva, Hernández-Ávalos et al. 2023). However, reference ranges are not very common in mice, often highly specific in their nature for the mouse model and typically based on very small numbers of mice (Rottman, Ni et al. 2007) (Stypmann, Engelen et al. 2006) (Vinhas, Araújo et al. 2013).

A recent position paper by the ESC Working Group on Myocardial Function outlines basic requirements and reference ranges for assessing echocardiography in mice and rats (Zacchigna, Paldino et al. 2021). To expand these existing resources and prove their validity, we leverage the extensive International Mouse Phenotyping Consortium (IMPC) resource and generate TTE reference ranges from over 15,000 conscious and anesthetized C57BL/6N wildtype control mice stratified by sex and age. Using this unprecedented scale of data we generated a generalizable framework for defining ‘normal’ morphology and functionality of the mouse heart.

## Materials/Methods

### The International Mouse Phenotyping Consortium

The International Mouse Phenotyping Consortium (IMPC) represents a multi-institutional and collaborative research initiative encompassing twenty-four major research organizations and funding agencies, distributed globally (Dickinson, Flenniken et al. 2016). The IMPC seeks to generate and phenotype a knockout mouse line for every protein-coding gene in the orthologous mouse genome (www.mousephenotype.org) (Muñoz-Fuentes, Cacheiro et al. 2018). Phenotyping is carried out under the uniform operating procedures detailed in IMPReSS (International Mouse Phenotyping Resource of Standardized Screens; www.mousephenotype.org/impress/index), which were developed and validated during the pilot programs EUMORPHIA and EUMODIC (Brown, Chambon et al. 2005).

### IMPC Centers Contributing Transthoracic Echocardiography Data

IMPC data release (DR) 21 was used herein (https://www.mousephenotype.org/data/release). The following subset of six IMPC data-contributing centers provided transthoracic echocardiography (TTE) data in DR 21 (ethical approval details are included in parenthesis after each contributing center):

1. Baylor College of Medicine (BCM) (Institutional Animal Care and Use Committee approved license AN-5896).
2. German Mouse Clinic, Helmholtz Zentrum München (GMC) (#144-10, 15-168)
3. Medical Research Council (MRC) – Harwell (HAR) (Animal Welfare and Ethical Review Body approved licenses 70/8015 and 30/3384).
4. Institute Clinique de la Souris, Mouse Clinical Institute (ICS) (#4789-2016040511578546v2).
5. Czech Centre for Phenogenomics (CCP) (AV CR 62/2016, Academy of Sci., Czech Rep.).
6. MARC Nanjing University (#NRCMM9).

TTE data was collected from mice at one of two possible time points. For the Early Adult (EA) Pipeline, data were collected at a mean of 12 weeks with the minimum of 8 and maximum of 16 weeks of age. For the Late Adult (LA) Pipeline, data were collected at a mean of 63 weeks with the minimum of 51 and maximum of 78 weeks of age. Animal welfare was assessed routinely for all mice involved.

### Animals

This study includes data collected from inbred wildtype control animals tested as part of the IMPC data. These mice, both males and females, were on a C57BL/6N genetic background of substrains: C57BL/6NCrl (CCP, HMGU and ICS); C57BL/6NJ (BCM) and C57BL/6NTac (HMGU, ICS and HAR).

Non-IMPC mice were from three different studies: (1) The founder strains animals from a study titled “The Collaborative Cross: A Recombinant Inbred Mouse Population for the Systems Genetic Era” (Threadgill, Miller et al. 2011) with A/J, C57BL/6J, 129S1/SvlmJ, NOD/ShiLtJ, NZO/HlLtJ, CAST/EiJ, PWK/Ph, and WSB/EiJ inbred strains (https://phenome.jax.org/projects/GMC13); (2) The Jaxwest1 project, a multi-system analysis of physiology on seven inbred strains of mice: 129S1/SvImJ, A/J, BALB/cJ, C57BL/6J, DBA/2J, NOD/ShiLtJ and SJL/J (https://phenome.jax.org/projects/Jaxwest1); (3) Eumorphia6, a project conducted under the Eumorphia / Europhenome initiative, is a collaborative effort involving multiple European research centers focused on phenotyping and genetic research on the inbred strains 129S2/SvPas, BALB/cByJ, C3HeB/FeJ and C57BL/6J (https://phenome.jax.org/projects/Eumorphia6).

### Transthoracic Echocardiogram (TTE) recording

The IMPC standard operating procedure provides an overview of the conscious and anesthetized TTE procedures used by contributing centers (https://www.mousephenotype.org/impress/ProcedureInfo?procID=109).

In brief, bodyweights were taken shortly before transthoracic echocardiography. For anesthetized TTE recordings, the animal was placed in an induction chamber and anesthetized with 1.5-3% isoflurane or injected with tribromoethanol as an injectable anesthetic. While sedated, either as part of the TTE session or as a separate preparatory procedure, the animal undergoes hair removal of the chest. With the hair removed, the animal was placed on the imaging platform with its paws taped to ECG surface electrodes and a rectal probe inserted to monitor body temperature which was maintained at 36-37°C. During imaging, anesthesia was adjusted to maintain proper heart rate and keep the animal from waking up.

For awake TTE examinations, the animal was firmly held by the nape (in the supine position) in the palm of one hand with the tail held tightly between the last two fingers.

To facilitate ultrasound imaging, pre-warmed ultrasound gel was placed on the chest at the area of imaging, and transthoracic echocardiography recordings captured. For short-axis mode papillary muscles were used as an anatomic point of reference. For parasternal long-axis, the apex and aortic root were used as anatomic points of reference. M-Mode images were captured, with at least 3 images per mode. Once imaging was complete, the animal was removed from the platform and allowed to recover atop a heating pad.

Measurements were made offline using analytical software (VisualSonics Inc.). Left ventricular diameter in systole (LVID;s) and diastole (LVID;d), as well as anterior and posterior wall thicknesses were measured for end-systolic and end-diastolic values (LVAW;s, LVPW;s, LVAW;d, LVPW;d. The papillary muscles, which serve as an anatomic marker for appropriate imaging position, were excluded from the traced boundary. Fractional shortening (FS) was calculated as FS % = [(LVID;d − LVID;s)/LVID;d] × 100. Ejection fraction (EF) was calculated as EF % = 100 × [(LVvol;d − LVvol;s)/LVvol;d] with LVvol = [(7.0/(2.4 + LVID) × LVID^3^]. The stroke volume (SV) is the volume of blood pumped from one ventricle of the heart with each beat. The stroke volume of the left ventricle was obtained by subtracting end-diastolic volume (LVvol;d) from end-systolic volume (LVvol;s). Heart rate was determined from the cardiac cycles recorded on the M-mode tracing, using at least three consecutive end-systolic intervals.

Importantly, each center records significant metadata parameters according to the FAIR principles (Findable, Accessible, Interoperable, Reusable; https://www.go-fair.org/fair-principles/) including equipment manufacturer; equipment model; recording environment; anesthetic agent; and anesthetic dose. Detailed experimental protocols on the IMPC phenotyping procedures are available for general access at www.mousephenotyping.org/IMPReSS. Data were curated and subject to quality control at the IMPC prior to Data Release (DR) 21.1 (13th June 2024) and we excluded four EA and one LA mice from the analysis due to body temperatures outside of a normal physiological range (<36°C, >40°C), two EA animals with improbable LVID;d < LVID;s values and two additional EA mice due to biologically implausible ejection fraction levels (<25%) in the conscious state.

## Statistical Methods

### Bespoke methods were developed to assess TTE reference ranges and are independent of the methodologies implemented on the IMPC portal

Data analysis was conducted using R (version 4.2.2, R Core Team 2022 (Team 2022) with figures and tables produced in ggplot2 and ggpubr. Variability of all the data was assessed by the metric coefficient of variation (COV). Visual methods (histograms and qqplots), as well as a formal statistical test (Shapiro-Wilks-test) were conducted to test whether the scores of the individual parameters were normally distributed. Data were separated by age, sex and anesthesia regime and histograms for each parameter were plotted. Reference ranges were calculated based on median, 25th percentile and 75th percentile. In addition, the mean, standard deviation, and parameter sample size were provided to reflect the distribution of each parameter. Based on this, the 95% confidence intervals can be calculated by mean±1.96*standard deviation for each parameter.

### Relative importance of predictors

To investigate the relative importance of different regressors on the outcome variable in linear models, we applied the R package “relaimpo” developed by Groemping (Groemping 2006) and calculated the relative importance (based on the metric ‘lmg’) of four predictors, namely anesthesia (conscious, isoflurane and tribromoethanol), sex (males and females), body weight and age (EA and LA) in all 15,765 mice. Adjusted R^2^ depicted the total proportion of variance explained by the model with all four predictors and the relative proportion of contribution for each predictor was shown by relative (%) of adjusted R^2^.

### Investigation of Anesthesia, Sex and Age Effects

To investigate the effect of anesthesia on the different parameters in EA mice, we calculated a one-way Analysis of Variance (ANOVA) with planned comparisons of “Conscious versus Isoflurane” and “Conscious versus Tribromoethanol”, separated by sex whereas “Isoflurane versus Tribromoethanol” was not tested. These planned comparisons were used to compare conscious vs unconscious states. The null hypothesis tested whether the two datasets originate from distributions with the same mean. P-values and F-values with degrees of freedom were calculated.

The effects of sex (female vs male) and age (EA vs LA) were compared using the identical statistical analyses. In each case a simple two-tailed t-test was performed and the Cohen’s d effect size calculated from the “effsize” package (R library). Due to the central limit theorem (CLT) (Zhang, Astivia et al. 2023), the large sample sizes allowed parametric statistical testing of these effects. The biological relevance of large samples may be overstated, which is why we also calculated the effect sizes in order to be able to estimate this factor.

## Data availability

All data used are available to the public for download at the IMPC (https://previous-releases-reports.s3.eu-west-2.amazonaws.com/release-21.pdf).

## Results

TTE data collected by IMPC contributing centers (data release, DR, 21) were available from 15,765 wildtype control mice, stratified as presented in Table 1 and summarized below. All the mice were from a C57BL/6N inbred substrain. TTE was performed on conscious mice, or mice anesthetized with either isoflurane or tribromoethanol. The majority of mice (89.3% or 14,083) were tested at a mean age of 12 weeks (designated as “early adult” or EA), while the remaining 10.7% (1,682) of mice were tested at a mean age of 63 weeks (designated as “late adult” or LA). Sex was evenly distributed at both EA and LA time points. Raw data can be downloaded using the following link: https://www.mousephenotype.org/data/previous-releases/21.1. The total number of reported parameters varied slightly between mice and can be accessed in each table.

**Table 1:** TTE data were available from a total of 15,765 mice, stratified by sex, age at testing (EA = 12 weeks of age; LA = mean of 63 weeks of age), and conscious state (conscious, anesthetized using isoflurane, or anesthetized using tribromoethanol).

|  | Conscious | Isoflurane | Tribromoethanol | Sum |
| --- | --- | --- | --- | --- |
| *EA female* | 2,699 | 4,633 | 227 | 7,559 |
| *EA male* | 2,709 | 3,592 | 223 | 6,524 |
| *LA female* | 378 | 496 | 0 | 874 |
| *LA male* | 376 | 432 | 0 | 808 |
| *Sum* | 6,162 | 9,153 | 450 | 15,765 |

## Variability Assessment

A panel of 12 output parameters were collected from TTE, namely cardiac output (CO), EF, LVID;d, LVID;s, FS, heart rate (HR), LVAW;d, LVAW;s, LVPWd, LVPWs, and SV (parameter definition in Supplemental Table 1). In addition, the body weight (BW) of each mouse is weighed before the TTE measurement, and the body temperature of anesthetized mice is measured with a rectal probe to monitor physiological functions (Supplemental Figure 1 with histograms, mean ±SD, median and 95% reference ranges, separated by sex). LVAW;d and LVAW;s were inconsistently collected in the IMPC and not further processed in this analysis.

In multi-center, large-scale, high-throughput programs such as the IMPC, variability in the measured values was to be expected. However, the extent of this variability dictates the sensitivity and robustness of each parameter. Variability testing was performed on all DR 21 TTE data from the IMPC, independently of anesthetic agent in this analysis. For each sex, individual TTE parameters were tested for variability in EA and LA populations. The coefficient of variation (COV) (100*standard deviation/mean) assumes a parametric distribution and normalizes the variability to the most typical score (mean) but is sensitive to outliers (2008). Based on this analysis, exclusion criteria were defined as any parameter with ≥30 for COV, based upon Eurachem guidelines (Williams 2012). Figure 1 shows that the retained parameters are all clustered closely together, however the excluded parameter shows a wide range of variability.

**Figure 1:**
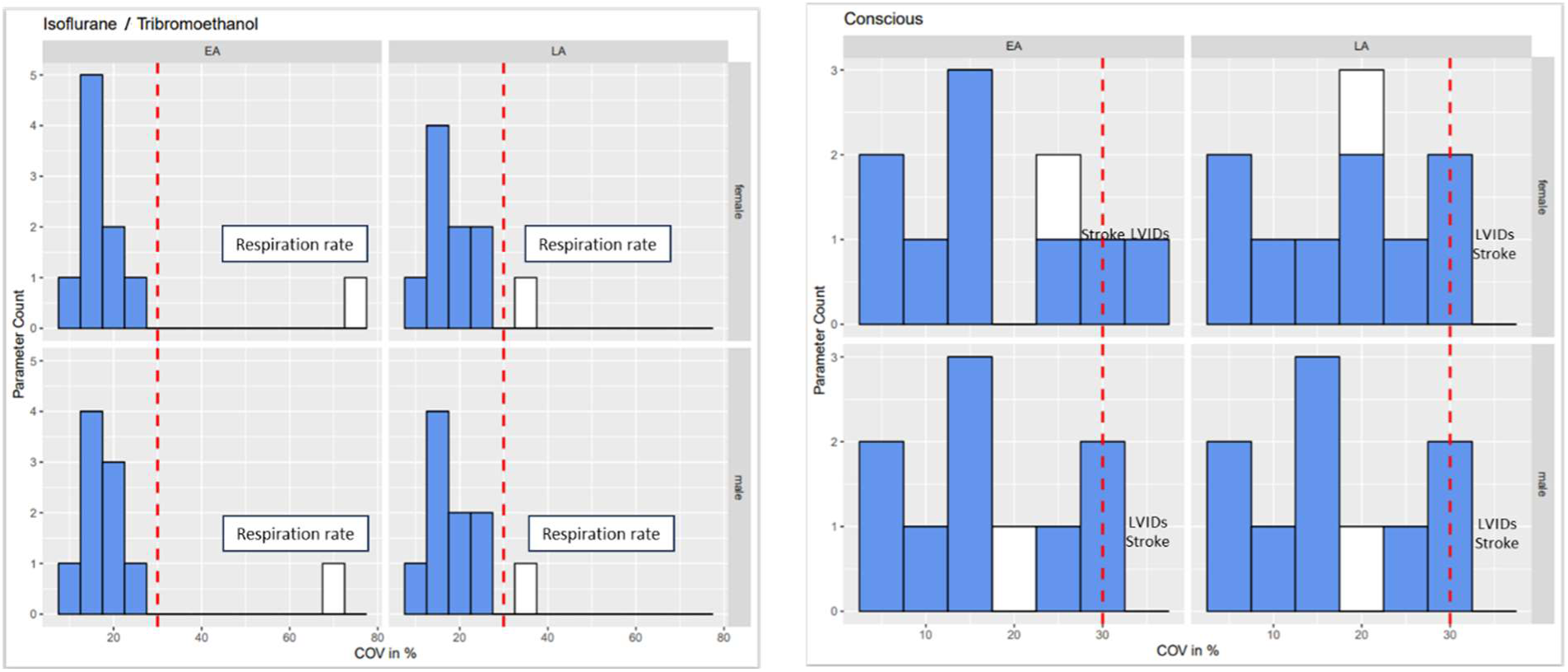
Coefficient of variation (COV) analysis of data, stratified by sex (female and male), anesthetic regime and age (EA and LA) identified one parameter, respiration rate, with excess variability (COV >30%) in LA males and females under anesthesia that was excluded from further analysis (white bar). Parameters in blue were below the COV threshold of 30% and were retained for further analysis.

Specifically, one TTE-parameter, respiration rate, exceeded the variability criteria in both sexes (male and female) and LA age but not in EA and was excluded from further analysis (Figure 1). The variability threshold was partially exceeded for LVID;s in females of EA age, however, in females of LA and males of both ages (EA and LA) the threshold was not exceeded and LVID;s was retained, whereas LVAW;d and LVAW;s values are exclusively shown in supplementals.

The remaining 9 TTE-parameters (CO, EF, LVID;d, LVID;s, FS, HR, LVPW;d, LVPW;s and SV) consistently presented with low variability across the whole IMPC dataset thereby giving high confidence to establish robust, generalizable reference ranges for EA and LA populations on the C57BL/6N inbred genetic background.

## Assessment of Data Distribution

The distribution of data was assessed via histograms for the nine selected TTE parameters stratified by sex, age, and anesthetic regime (Figure 2). Under tribromoethanol anesthesia, data points were only captured in EA from five (FS, HR, LVID;d, LVID;s and LVPW;d) of the nine TTE parameters.

**Figure 2:**
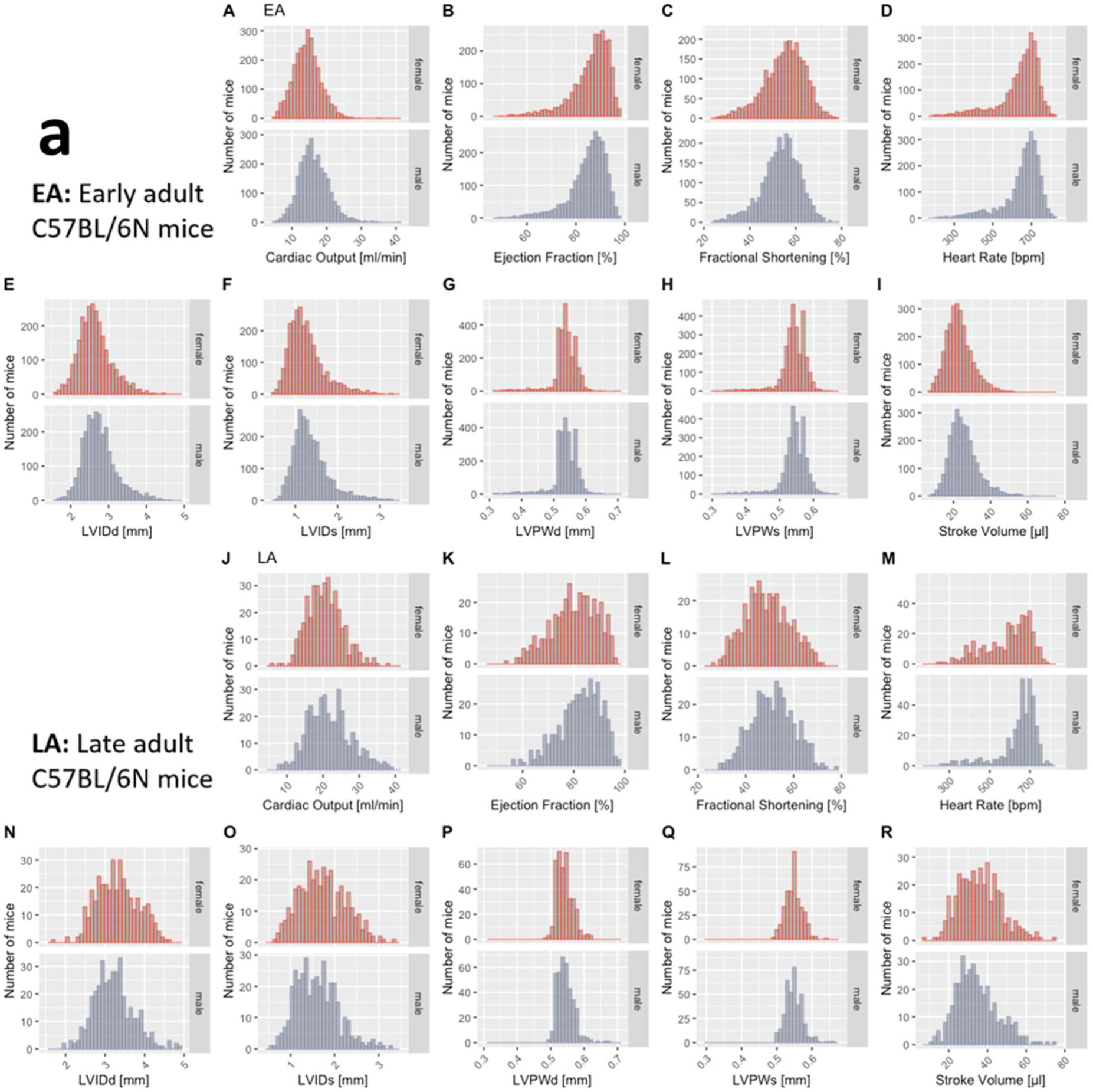
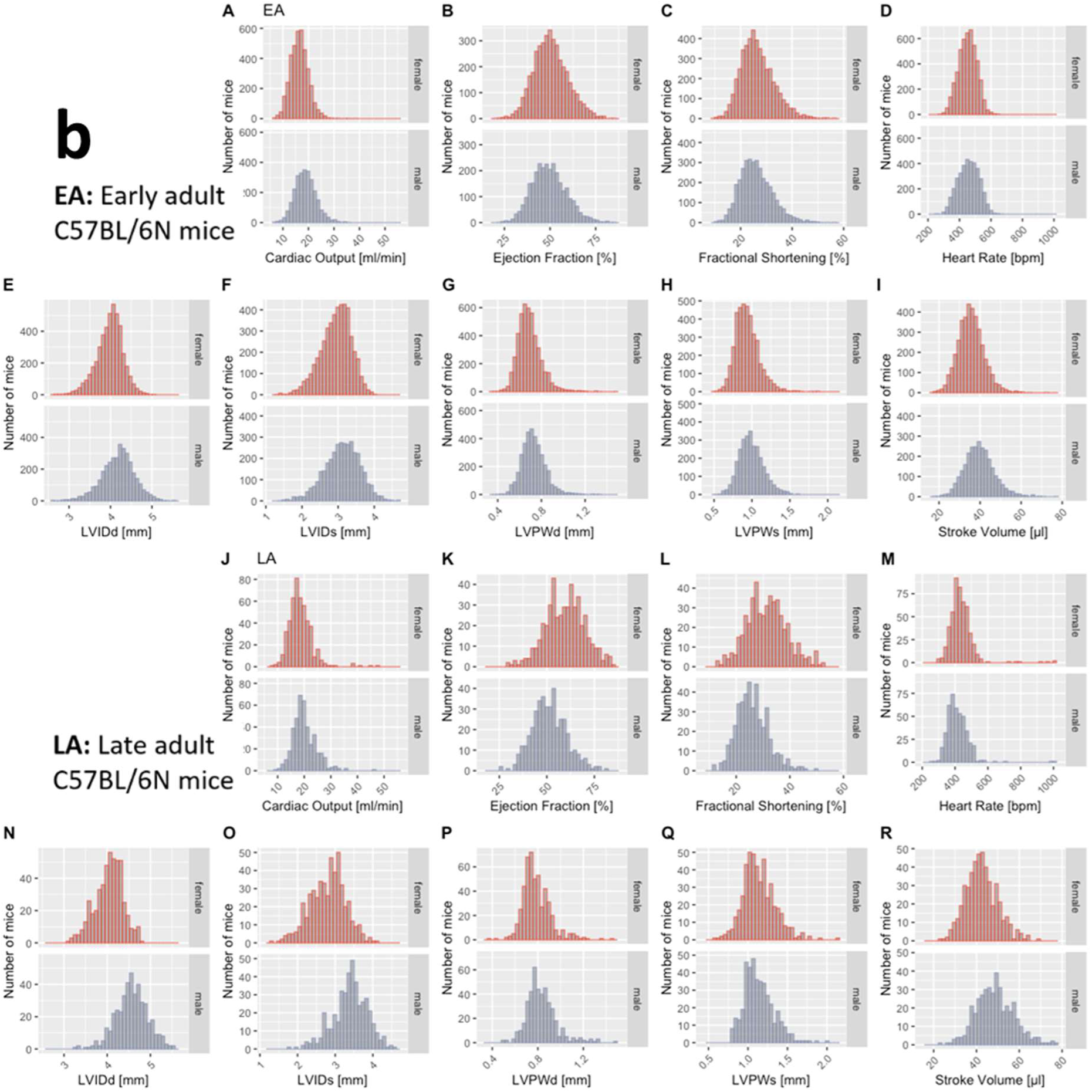
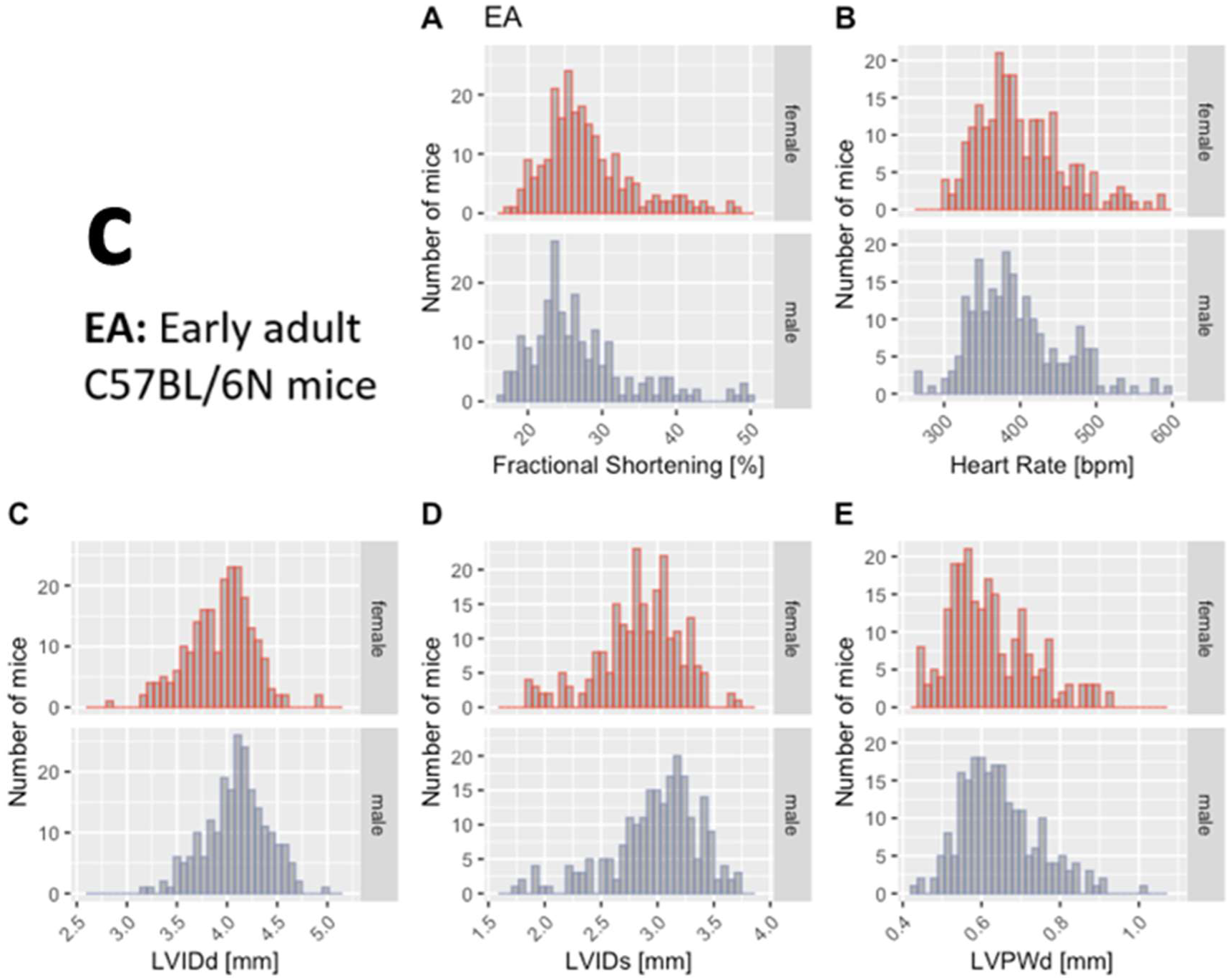
Histograms presenting the distribution of each selected TTE parameter for male (blue) and female (red) mice separately. By visual inspection, no sexual dimorphism is apparent. Panel a: Recorded in the conscious state in early adult, EA (Sub-panels A-I) and late adult (LA) mice (Sub-panels J-R). Panel b: Recorded under isoflurane anesthesia in early adult, EA (Sub-panels A-I) and late adult (LA) mice (Sub-panels J-R). Panel c: Recorded under tribromoethanol anesthesia in early adult, EA (Sub-panels A-E) with five (FS, HR, LVID;d, LVID;s and LVPW;d) out of nine TTE parameters. No late adult data are available for tribromoethanol anesthesia.

This visual representation of the frequency of occurrence per value in the data was useful for revealing conformity to and deviations from a normal distribution for each parameter. Visual inspection of the histograms showed that the data appeared practically normal in EA for parameters CO, EF, LVID;d, LVID;s, FS, LVPW;d, LVPW;s and SV and modestly skewed for HR in conscious mice but not under isoflurane anesthesia. Under tribromoethanol anesthesia, FS, HR, LVID;d, LVID;s and LVPW;d appeared practically normally distributed whereas HR was modestly skewed. To assess normality mathematically, we applied the Shapiro-Wilk test which revealed statistically significant deviation from a normal distribution for some, but not all, TTE parameters. Table 2 presents data as median and 95% reference range (2.5th and 97.5th percentile) to account for the lack of normal distribution of some parameters and to provide a consistent data presentation. Data are stratified by sex, age (EA and LA) and conscious state. For the sake of completeness, mean, standard deviation and sample size are provided for the seven selected TTE parameters stratified by sex, age, and anesthetic regime in Supplemental Table 2 and LVAW;d and LVAW;s values in Supplemental Table 3.

**Table 2:** Median and 95% reference ranges of CO, EF, FS, HR, LVID;d, LVID;s, LVPW;d, LVPW;s, and SV. Data are stratified by sex, age (EA and LA) and conscious state. Panel a. Conscious, isoflurane and tribromoethanol anesthetized female mice. Panel b. Conscious, isoflurane and tribromoethanol anesthetized male mice. Note, there were no data for LA mice anesthetized using tribromoethanol.

| **a** | Conscious - Females |  | Isoflurane - Females |  | Tribromoethanol - Females |  |
| --- | --- | --- | --- | --- | --- | --- |
|  | EA | LA | EA | LA | EA | LA |
| *Parameter* | median [95% range] | median [95% range] | median [95% range] | median [95% range] | median [95% range] | no data available |
| *Cardiac Output [ml/min]* | 14.3 [7;22.5] | 20.3 [12.3;31.7] | 16.4 [10.1;23.8] | 17.9 [12.1;27.1] | no data available |  |
| *Ejection Fraction [%]* | 88 [65;96] | 81.1 [60.6;94.7] | 49.9 [33.9;70.3] | 59.9 [38.6;81.2] | no data available |  |
| *Fractional Shortening [%]* | 56.1 [34.9;70.1] | 48.8 [32.2;67.3] | 25.8 [16.2;40.7] | 31.2 [17.5;48.6] | 26.7 [19.6;43.1] |  |
| *Heart Rate [bpm]* | 668.2 [336.8;769.2] | 620.2 [343;751.3] | 454 [342;556.3] | 427 [339.4;549] | 390 [315.2;538.7] |  |
| *LVIDd [mm]* | 2.6 [1.9;3.6] | 3.3 [2.5;4.3] | 4 [3.3;4.6] | 4.1 [3.4;4.7] | 4 [3.2;4.5] |  |
| *LVIDs [mm]* | 1.2 [0.7;2.4] | 1.7 [0.8;2.8] | 3 [2;3.7] | 2.9 [1.8;3.7] | 2.9 [1.9;3.4] |  |
| *LVPWd [mm]* | 0.5 [0.4;0.6] | 0.5 [0.5;0.6] | 0.7 [0.5;0.9] | 0.8 [0.6;1.1] | 0.6 [0.4;0.9] |  |
| *LVPWs [mm]* | 0.5 [0.4;0.6] | 0.6 [0.5;0.6] | 0.9 [0.7;1.3] | 1.1 [0.8;1.6] | no data available |  |
| *Stroke Volume [μl]* | 22.3 [11;39.7] | 35.7 [18.3;59.6] | 35.3 [24.3;48.8] | 41.9 [29.8;58.9] | no data available |  |

## Relative importance of predictors

If there are several predictors, the question naturally arises as to which predictor is more important or useful for predicting the outcome variable. For correlated predictors, the standardized coefficients may not indicate which predictor is more important. Here, the calculation of the ‘relative importance’ of the predictors (Groemping 2006) is applied for the predictors sex (males and females), anesthesia (conscious, isoflurane and tribromoethanol), age (EA and LA) and body weight across the grand total of 15,765 mice. Anesthesia showed highest proportion of contribution with a range of 32 (SV) to 99% (EF and FS) relative % of adjusted R^2^ for all 9 TTE parameters wherein sex, weight and age covered rather low proportions of variance (Table 3).

**Table 3:** Adjusted R^2^ represents the total explained variance of the relaimpo model with the proportion of contribution for each predictor shown by relative % of adjusted R^2^ for anesthesia (conscious, isoflurane or tribromoethanol anesthesia), sex (female and male), (body) weight, and age (EA and LA) across the total of 15,765 mice. Anesthesia contributes with a range of 32 to 99% to the adjusted R^2^ of all 9 TTE parameters.

| All TTE data from the IMPC |  |  |  |  |  |
| --- | --- | --- | --- | --- | --- |
|  | Adj. R<sup>2</sup><br>explained model<br>variance | Relative % of adj. R<sup>2</sup> |  |  |  |
|  |  | Anaesthesia | Sex | Weight | Age |
| Cardiac Output | **18%** | 32% | 16% | 39% | 13% |
| Ejection Fraction | **81%** | 99% | 0.3% | 0.3% | 0.3% |
| Fractional Shortening | **80%** | 99% | 0.3% | 0.3% | 0.3% |
| Heart Rate | **62%** | 96% | 1% | 1% | 2% |
| LVIDd | **80%** | 95% | 0% | 3% | 2% |
| LVIDs | **82%** | 98% | 0% | 1% | 1% |
| LVPWd | **52%** | 89% | 1% | 5% | 5% |
| LVPWs | **75%** | 94% | 0% | 2% | 4% |
| Stroke Volume | **56%** | 78% | 2% | 12% | 8% |

## Effect of Sex

Interestingly, male, and female data showed similar distributions by visual inspection (Figure 2). To test the hypothesis that there is no difference between each sex, a simple two-tailed t-test was performed independently for each anesthetic regime and age group, and Cohen’s d was calculated as an effect size measure (Supplemental Figure 2, 3 and 4 – Panels a and b, stratified by age). In the EA population, for some parameters, p-values reached significance <.001, for others we found no evidence of a difference (Panel a). However, for all parameters the corresponding Cohen’s d value revealed small to negligible effect sizes. We therefore considered the possibility that the large group sizes could be overstating the biological differences between the sexes for some parameters. In the LA population, p-values reached significance <.001 for most of the parameters and for others we found no evidence of a difference (Panel b). In contrast, the corresponding Cohen’s d value revealed small to negligible effect sizes for the majority of parameters, however, large effect sizes were observed in EF, LVIDd, LVIDs and FS in mice under isoflurane and medium for SV. Here, with a relatively smaller group size than EA population, the biological differences between the sexes in the LA population are both significant and of medium or large effect sizes for some, mainly functional (and less morphological), TTE parameters. Overall, there is a difference between sexes.

## Effect of Anesthetic Agent

To investigate the effect of different anesthetic agents on cardiac conduction function and TTE profiles, conscious data stratified by sex and age are displayed for comparison with those of isoflurane or tribromoethanol data (Figure 3). Female data are placed directly above male for ease of visualization. Figure 4 shows distinct distribution clusters for conscious, isoflurane and tribromoethanol groups split by EA (Figure 4 – Panels A to G) and LA (Figure 4 – Panels H to N). As before, no data were available for tribromoethanol anesthesia in LA mice.

**Figure 3:**
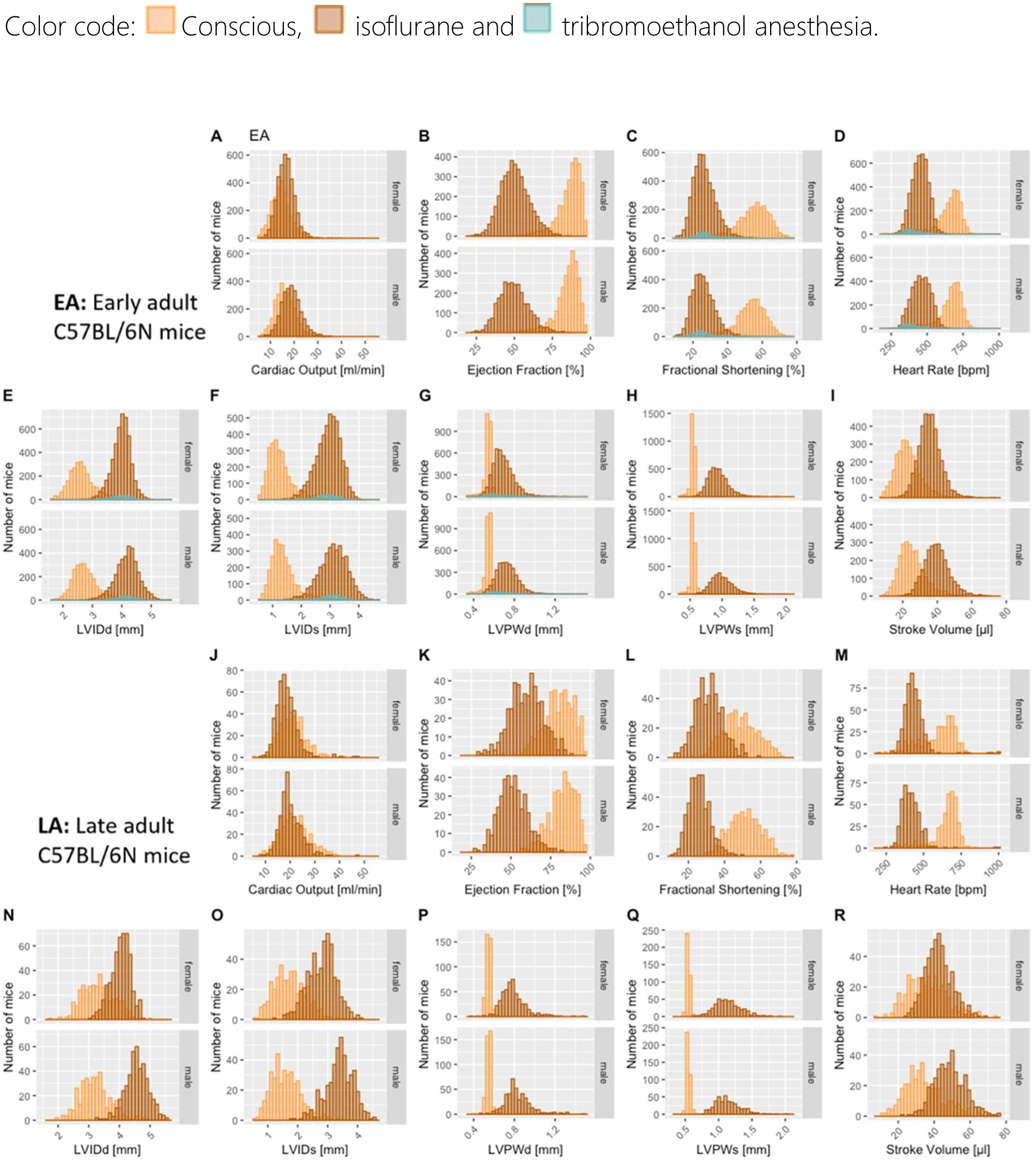
Comparison of the anesthetic regimes with the conscious state recordings. Distribution of the nine selected TTE parameters presented by histograms, stratified for female and male mice in early adult, EA (Sub-panels A-I) and late adult (LA) populations (Sub-panels J and R). No late adult data and only FS, HR, LVID;d, LVID;s and LVPW;d parameters in EA are available for tribromoethanol anesthesia.

**Figure 4:**
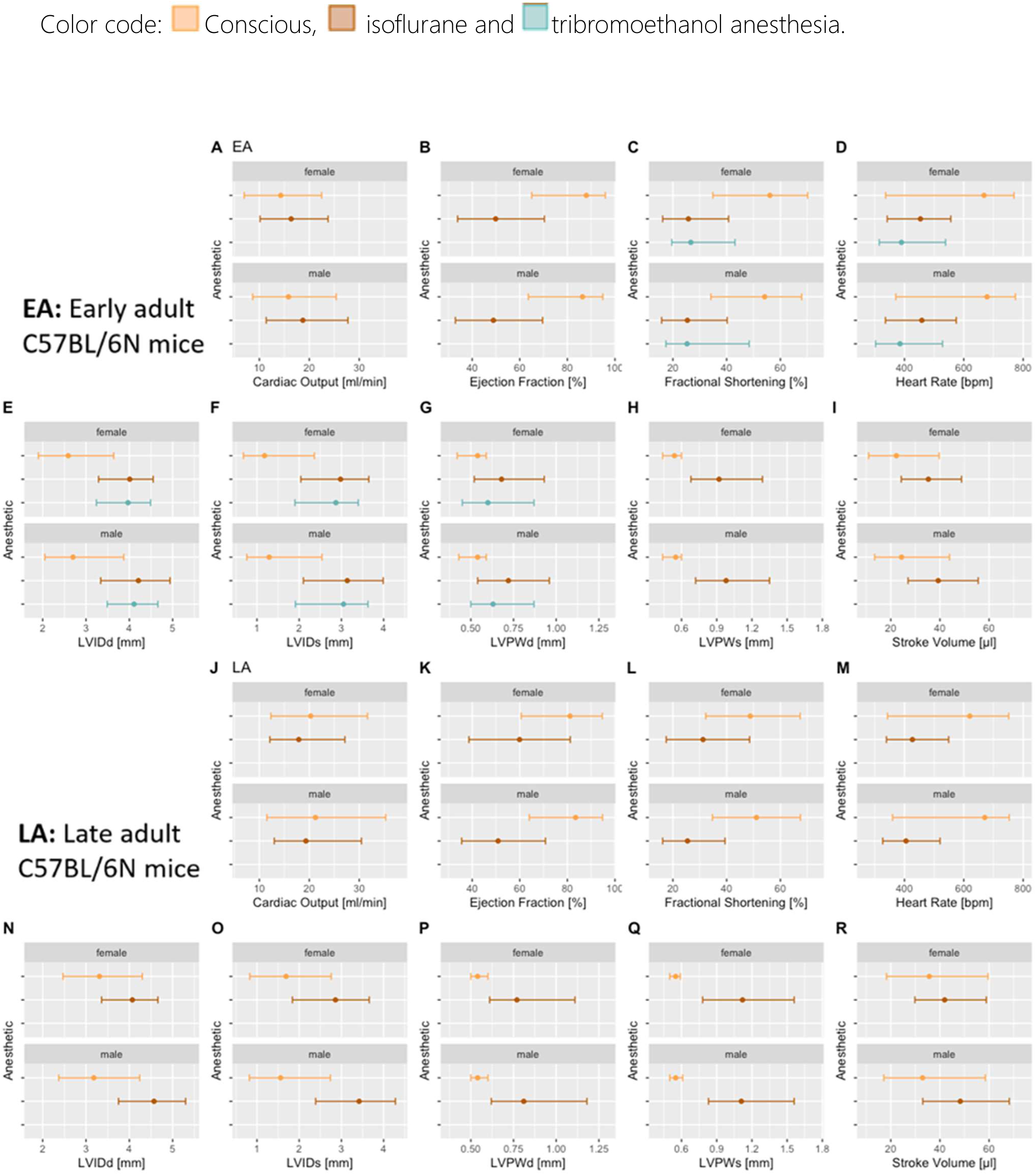
Reference ranges split by anesthetic regimen showing median, and 95% reference ranges (2.5th and 97.5th percentile). Female data are directly above the male data for EA (Sub-panels A-I) and late adult (LA) populations (Sub-panels J-R). No late adult data and only FS, HR, LVID;d, LVID;s and LVPW;d parameters in EA are available for tribromoethanol anesthesia.

As expected, the physiological benchmark of highest heart rate in conscious mice compared to anesthetized animals was observed in EA and LA populations (Figure 3 – Panels D and M). To assess the differences between EA anesthetic states, we tested conscious versus isoflurane and conscious versus tribromoethanol groups, by a one-way ANOVA with planned comparisons, and observed highly significant differences between those groups except for LVPWd in EA females conscious versus Tribromoethanol (Table 2). These data clearly show differences in TTE parameters that can be attributed to the anesthetic regime; therefore, it is essential to establish reference ranges separately by condition (conscious or anaesthetized) and by anesthetic (isoflurane or tribromoethanol).

## Effect of Age

Two different age groups, i.e. mean of 12-weeks (minimum 8 and maximum 16 weeks) old EA and mean of 63 weeks (minimum 51 and maximum 78 weeks) old LA, have made it possible to explore the effect of age on TTE parameters in conscious and isoflurane anaesthetized mice, stratified by sex. A two-tailed t-test was applied to test the difference between the means of EA and LA results in conscious (Table 4) and isoflurane anaesthetized mice (Table 5). P-values <.001 were reached for all parameters except HR and LVPW;s in conscious males, indicating high statistical significance and the corresponding Cohen’s d effect size revealed medium to large standardized effect sizes in CO, LVID;d, LVID;s and SV also EF and FS in conscious females (Table 4 – Panels a and b). In isoflurane anaesthetized mice, p-values <.001 were reached for all parameters except EF in males, indicating high statistical significance and the corresponding Cohen’s d effect size revealed medium to large standardized effect sizes in, LVID;d, LVID;s, LVPW;s and SV also CO, EF, FS and LVPW;d in females (Table 5 – Panels a and b). These high significance values with considerable effect sizes suggest that the influence of age strongly influenced the results independent of the anesthetic regime.

**Table 4:** Significant differences between the statistical comparison of conscious versus isoflurane (p<.001) and conscious versus tribromoethanol (p<.001 to p=.004) in female (Panel a) and male (Panel b) mice for CO, EF, FS, HR, LVID;d, LVID;s, LVPW;d, LVPW;s, and SV. Test: p-value and F-value of one-way ANOVA with planned comparison.

| Females |  |  |
| --- | --- | --- |
|  | Conscious vs Isoflurane | Conscious vs Tribromoethanol |
| Cardiac Output [ml/min] | F(1,6432)=527.2, $p < .001$ | no data available |
| Ejection Fraction [%] | F(1,6593)=26877.2, $p < .001$ | no data available |
| Fractional Shortening [%] | F(1,7538)=25534, $p < .001$ | F(1,7538)=683, $p < .001$ |
| Heart Rate [bpm] | F(1,7533)=9561.5, $p < .001$ | F(1,7533)=749.8, $p < .001$ |
| LVIDd [mm] | F(1,7536)=23387.1, $p < .001$ | F(1,7536)=648.2, $p < .001$ |
| LVIDs [mm] | F(1,7537)=27345.1, $p < .001$ | F(1,7537)=690, $p < .001$ |
| LVPWd [mm] | F(1,7537)=4984.7, $p < .001$ | F(1,7537)=0.7, $p = .42$ |
| LVPWs [mm] | F(1,6592)=16533.3, $p < .001$ | no data available |
| Stroke Volume [μl] | F(1,6454)=5677.1, $p < .001$ | no data available |

| Males |  |  |
| --- | --- | --- |
|  | Conscious vs Isoflurane | Conscious vs Tribromoethanol |
| Cardiac Output [ml/min] | F(1,5368)=512.6, $p < .001$ | no data available |
| Ejection Fraction [%] | F(1,5533)=23496.7, $p < .001$ | no data available |
| Fractional Shortening [%] | F(1,6504)=22155.6, $p < .001$ | F(1,6504)=668.7, $p < .001$ |
| Heart Rate [bpm] | F(1,6502)=9073, $p < .001$ | F(1,6502)=873.4, $p < .001$ |
| LVIDd [mm] | F(1,6502)=18163.3, $p < .001$ | F(1,6502)=501, $p < .001$ |
| LVIDs [mm] | F(1,6505)=21830.1, $p < .001$ | F(1,6505)=574, $p < .001$ |
| LVPWd [mm] | F(1,6504)=7485.5, $p < .001$ | F(1,6504)=5.1, $p = .024$ |
| LVPWs [mm] | F(1,5534)=19865, $p < .001$ | no data available |
| Stroke Volume [μl] | F(1,5381)=4961.7, $p < .001$ | no data available |

In summary, Figure 5 is a graphical violin representation of the median and the full distribution of the numeric data including the density of each TTE variable broken down by anesthetic regimen with the female data placed directly above equivalent male data for easy visual interpretation; corresponding numeric values are presented in Table 2. This graphical representation clearly shows that age has a strong influence with respect to the distribution and the reference ranges of nine TTE parameters.

**Figure 5:**
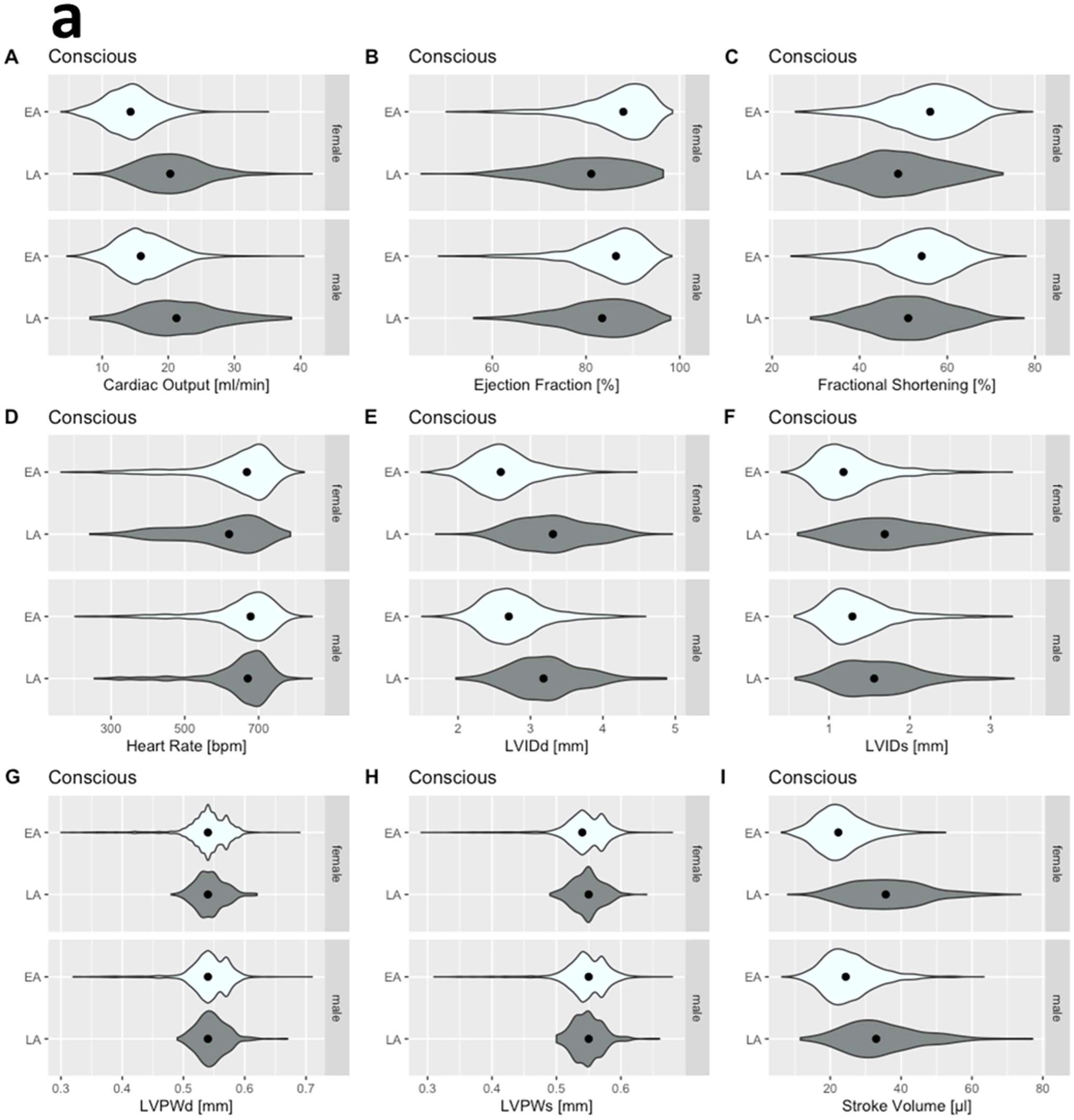
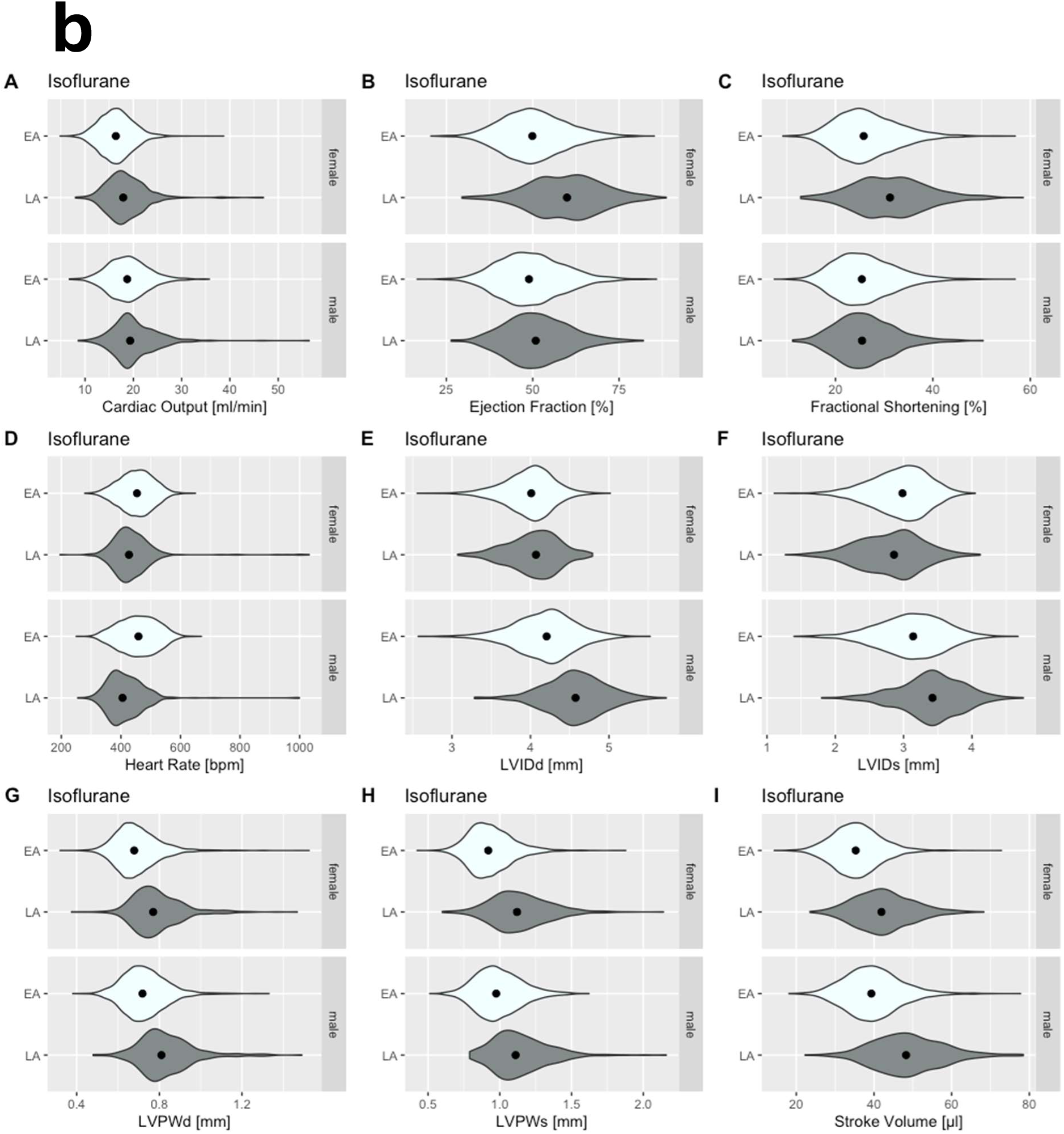
Violin plot representing full distribution of the numeric data including the density of each TTE variable. Female data are directly above the male data for late adult (LA) and early adult (EA) populations for easy comparison. Panel a: Recorded in the conscious state in EA and LA, Panel b recorded under isoflurane anesthesia. No late adult data are available for tribromoethanol anesthesia.

## Effect of Body Weight

In tandem with age, body weight (BW) can also have an influence on the heart and prompted the question of how old, yet healthy C57BL/6N mice change and modify TTE profiles. Two different age groups, i.e. mean of 12-weeks (minimum 8 and maximum 16 weeks) old EA and mean of 63 weeks (minimum 51 and maximum 78 weeks) old LA, have made it possible to explore the BW in conscious and isoflurane anaesthetized mice, stratified by sex. A two-tailed t-test was applied to test the difference between the means of EA and LA results in conscious (Figure 6 – Panel a) and isoflurane anaesthetized mice (Figure 6 – Panel b). P-values <.001 were reached for BW in EA versus LA, indicating high statistical significance between these two age groups independent of sex and anesthesia.

**Figure 6:**
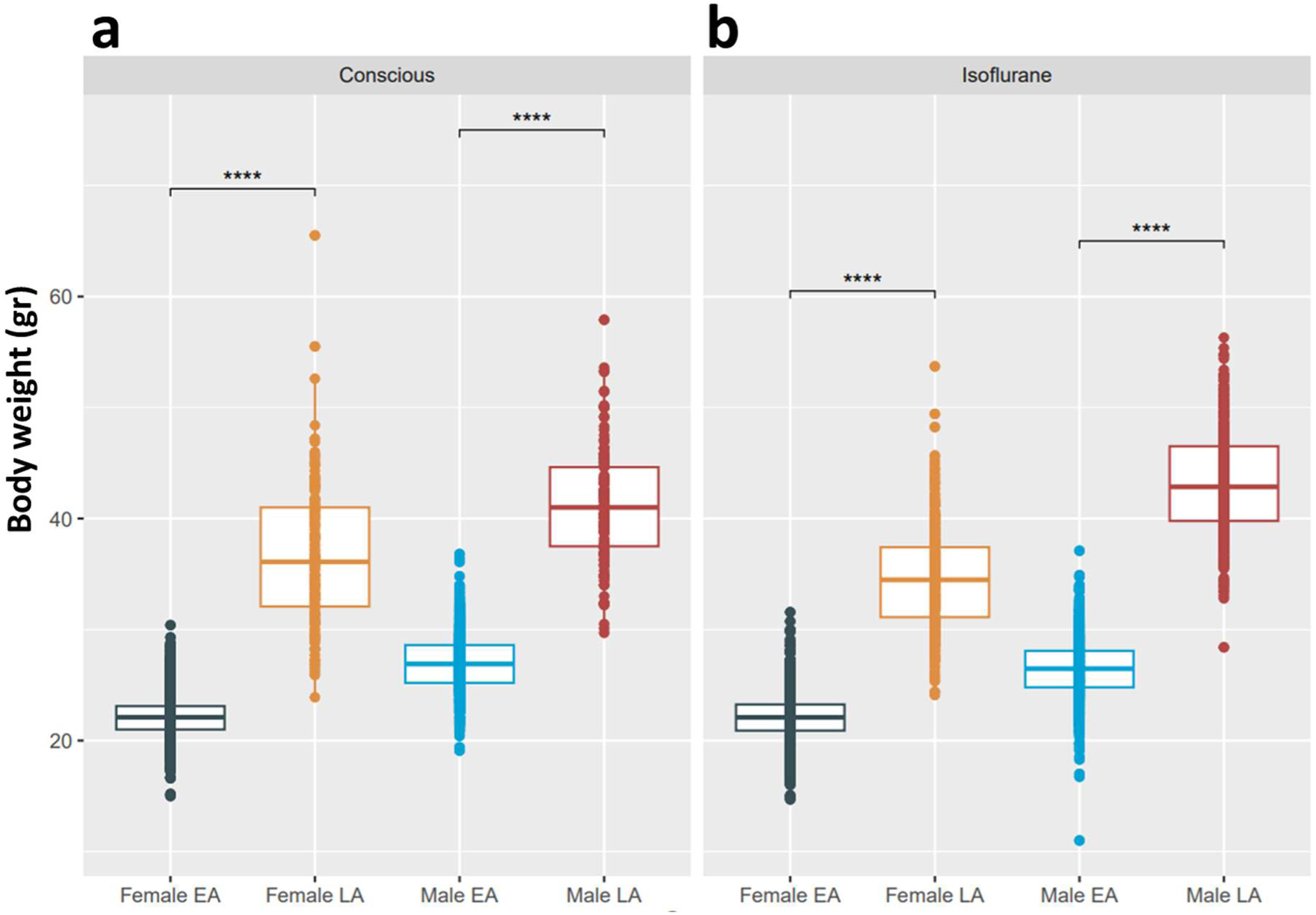
A two-tailed t-test was applied to test the difference between the means of EA and LA body weights (BW), stratified for female and male mice in early adult (EA) and late adult (LA) populations split by anesthetic regimen. Panel a: Recorded in the conscious state in EA and LA, Panel b recorded under isoflurane anesthesia. No BW data were available for tribromoethanol anesthesia. P-values <.001 (***) were reached for all comparisons, indicating high statistical significance between age groups.

To explore the linear relationship of BW and each TTE parameter, we calculated the Pearson correlation coefficient (Pearson’s R (Pearson 1895)), stratified by sex, age and anesthesia. Strength of the linear relationship between TTE variables and BW were defined based on user guidelines (Akoglu 2018, Schober, Boer et al. 2018): 0.00-0.10 negligible, 0.10-0.39 weak, 0.40-0.69 moderate, 0.70-0.89 strong and 0.90-1.00 very strong correlations. Female data are placed directly above male for ease of visualization.

Figure 7 shows distinct distribution clusters with regression line for conscious and isoflurane groups stratified by sex and split by EA and LA. No BW data were available for tribromoethanol anesthesia in EA and LA mice. Negligible to weak standardized Pearson correlation were reached for the entire set of TTE parameters with BW (gr) in conscious and isoflurane anesthetized mice ranging from 0.03-0.21 in the EA (Figure 7 – Panel a) and 0.01-0.38 in LA population (Figure 7– Panel b), indicating a minor role of BW; therefore, it is not entirely necessary to consider BW herein as major effect contributor.

**Figure 7:**
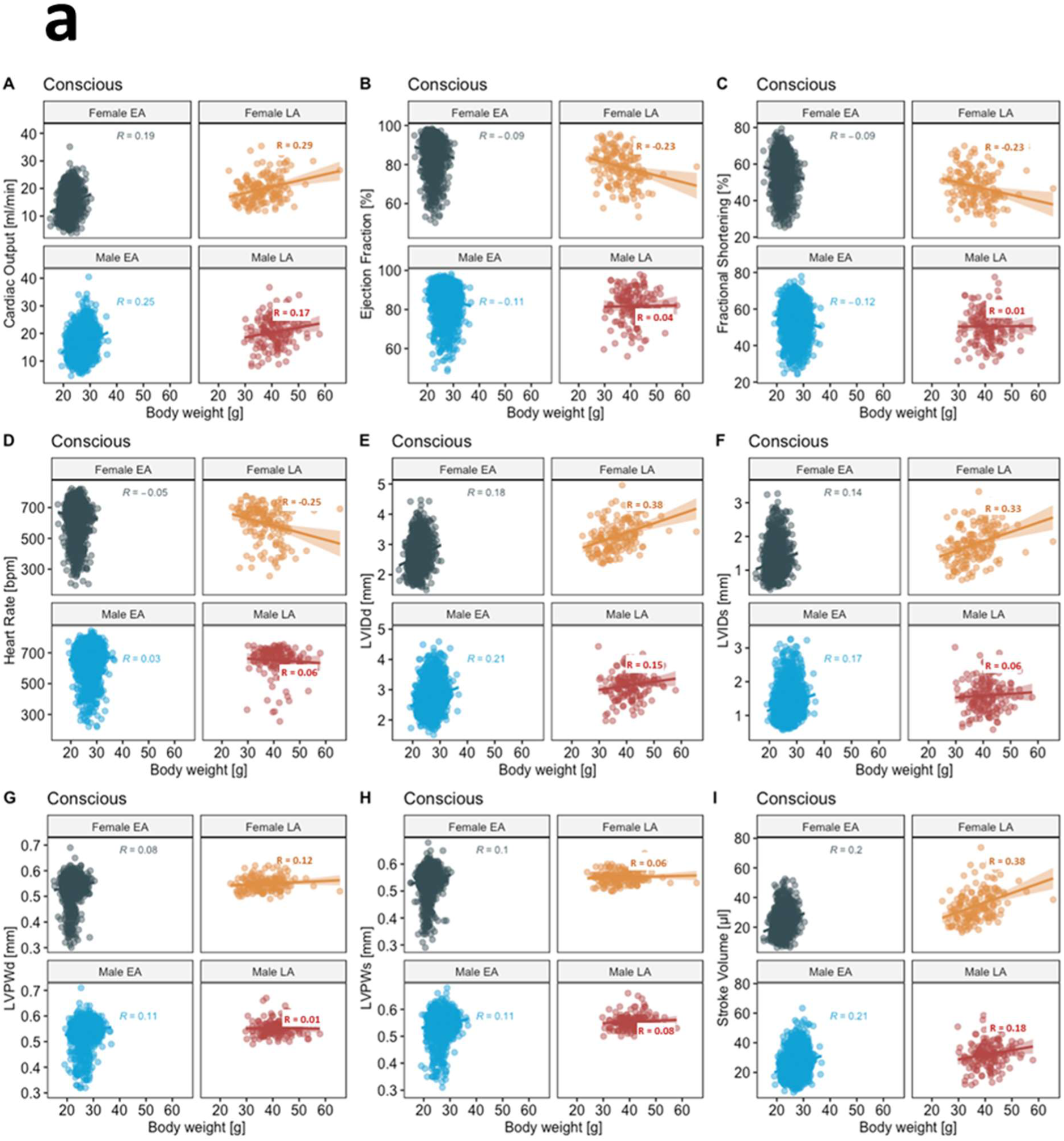
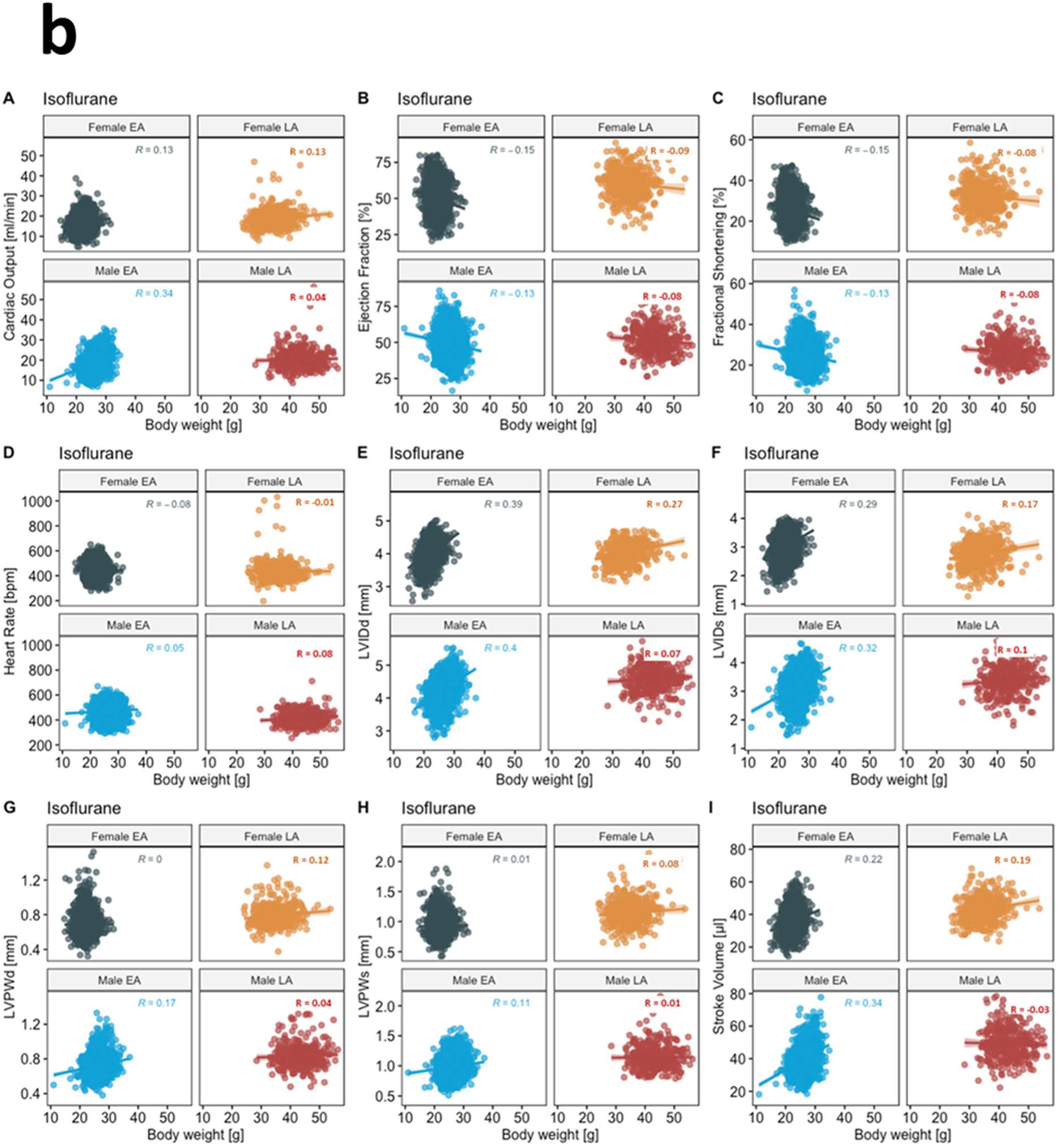
Pearson correlation with linear regression line presenting negligible to weak correlations for all nine TTE parameters with BW (gr), stratified for female and male mice in early adult, EA and late adult (LA) populations split by anesthetic regimen. Panel a: Recorded in the conscious state in EA and LA, Panel b recorded under isoflurane anesthesia. No late adult data are available for tribromoethanol anesthesia.

## Validation of Reference Ranges Using Non-IMPC Data

Mice characterized by the IMPC are all substrains of one commonly used inbred genetic background, C57BL/6N. To test the validity of the reference ranges reported herein beyond C57BL/6N inbred mice, we used data from representative additional inbred mouse strains from publicly available TTE data including: six founder strains from a collaborative cross study (Threadgill, Miller et al. 2011); the Jaxwest1 project (https://phenome.jax.org/projects/Jaxwest1) with seven inbred strains of mice; and four inbred strains of the Eumorphia / Europhenome project (https://phenome.jax.org/projects/Eumorphia6). An additional dataset was included using inbred, wildtype control animals from non-IMPC studies conducted at the German Mouse Clinic (GMC) where data is available upon request. Furthermore, we used published TTE reference ranges from the ESC position paper focusing on the appropriate echocardiographic acquisition and analysis of left ventricular function in healthy adult mice (Zacchigna, Paldino et al. 2021).

In each non-IMPC study, where suitable, we presented the data split by sex and overlaid with the sex-specific 95% reference range calculated herein for conscious (collaborative cross study and non-IMPC studies at GMC) or isoflurane anesthetized mice (Jaxwest1, Eumorphia6 and ESC study). In the ESC study, we used mean ± 1.96*SD to calculate 95% reference range, stratified by HR <450 b.p.m or HR >450 b.p.m, and compared them with the 95% reference range calculated herein, equally stratified according to the method of the ESC study. Figure 8 shows the founder strain data from the collaborative cross study overlaid with the reference ranges split by sex whereas Supplemental Figure 5 illustrates data from the Jaxwest1, Supplemental Figure 6 from Eumorphia6 and Supplemental Figure 7 for non-IMPC studies conducted at the German Mouse Clinic. Remarkably, and true for all TTE parameters, most non-C57BL/6N values lay within our reference ranges. There is a subset of outliers that fall outside of the reference ranges which is to be expected with heterogeneity of small size and phenotypic differences seen between inbred mouse strains. Figure 9 shows the ESC reference ranges overlaid with the reference ranges calculated herein, unclassified for sex. Remarkably, and true for all TTE parameters, most non-C57BL/6N values lay within our reference ranges. This visual representation of the reference ranges per value in the data was useful for revealing conformity to– and deviations from the reference ranges calculated herein, for each parameter. In both sets, HR <450 b.p.m or HR >450 b.p.m, there are minor shifts that fall beyond the reference ranges generated herein, which is to be expected with the heterogeneity of the ESC multicenter study of comparatively small mouse numbers and phenotypic differences between inbred mouse strains.

**Figure 8:**
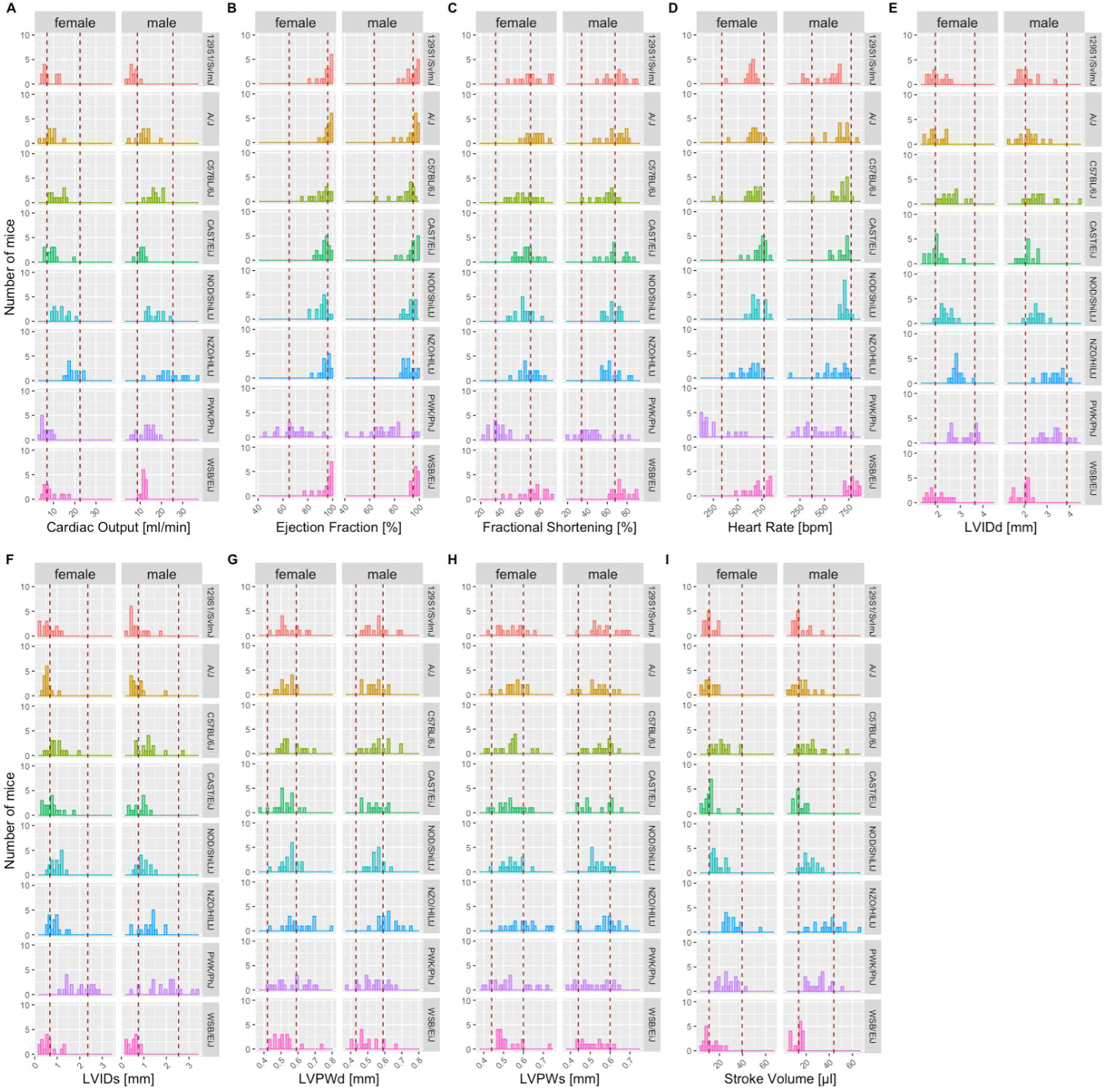
Independent, non-IMPC study on six of the founder strain mice reported in “The Collaborative Cross: A Recombinant Inbred Mouse Population for the Systems Genetic Era”(Threadgill, Miller et al. 2011) study, including 129S1/SvlmJ, A/J, C57BL/6J, NOD/ShiLtJ, NZO/HlLtJ, and PWK/PhJ inbred strains show a close alignment to the reference ranges reported herein for CO, EF, LVIDd, LVIDs, FS, HR, LVPWd, LVPWs and SV based on multiple C57BL/6N substrains indicating good utility for those reference ranges. Mice were conscious, split by sex and ∼12 weeks of age, equivalent to the IMPC EA time point. Red dotted lines depict the boundaries of the sex-specific reference ranges calculated from the C57BL/6N IMPC animals described earlier, for each parameter.

**Figure 9:**
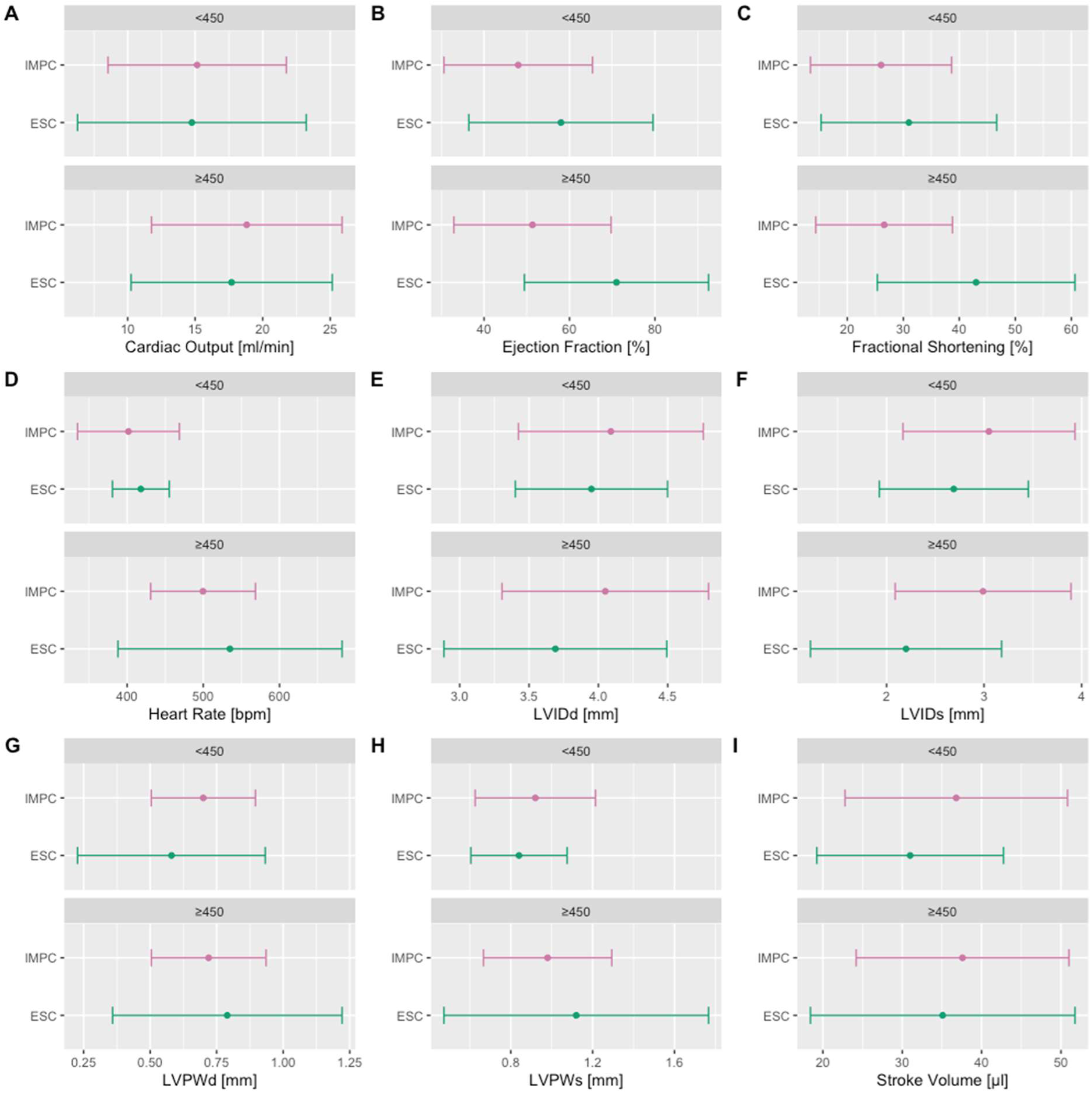
Independent, non-IMPC reference ranges of the ESC multicenter study including a total of 300 TTE exams, stratified by HR <450 b.p.m or HR >450 b.p.m, performed on control/healthy C57BL/6 adult mice across ten different laboratories reported in “Towards standardization of echocardiography for the evaluation of left ventricular function in adult rodents: a position paper of the ESC Working Group on Myocardial Function” (Zacchigna, Paldino et al. 2021). IMPC-mice were under isoflurane anesthesia whereas ESC-mice were under isoflurane or sevoflurane anesthesia. ESC-mice were pooled for sex and ∼12 weeks of age, equivalent to the IMPC EA time point. In both sets, <450 b.p.m or >450 b.p.m, CO, HR, LVPW;d, LVPW;s and SV show a close alignment to the reference ranges generated herein. There are minor shifts that fall beyond the reference ranges generated herein for EF and FS and concurrently LVID;d and LVID;s. Color code: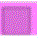 Reference ranges generated herein, 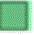 reference ranges of the ESC study, pooled for sex and stratified by <450 b.p.m (upper panel) or >450 b.p.m (lower panel).

## Discussion

Transthoracic echocardiography (TTE) is the imaging modality most widely used in cardiology (Galderisi, Cosyns et al. 2017). While TTE is non-invasive, it plays an important role in the diagnosis of numerous cardiovascular diseases providing valuable qualitative and quantitative insights into prognosis and pathophysiological processes (Boyd, Schiller et al. 2015). The European Association of Cardiovascular Imaging and the American Heart Association embody an acknowledged expert consensus in cardiology establishing, regularly revised, guidelines for the assessment and evaluation of echocardiography. (Galderisi, Cosyns et al. 2017, Douglas, Carabello et al. 2019). The availability of normal reference intervals for comparing individual patients is a cornerstone of echocardiography (Heidenreich 2023). Recent years have seen the publication of large, international, prospectively recruited studies with respect to age, gender and ethnicity, expanding towards risk assessments (Cain, Ahl et al. 2009, Jasaityte, D’Hooge et al. 2013, Lancellotti, Badano et al. 2013, 2014, 2015, Harkness, Ring et al. 2020, Wehner, Jing et al. 2020, Kondo, Dewan et al. 2023, Nyberg, Jakobsen et al. 2023).

Unlike other animals, mouse-specific reference ranges are uncommon (Konrad, Weber et al. 2000, Grenacher and Schwarzwald 2010, Vernemmen, Vera et al. 2020, Cerbu, Cerbu et al. 2023, Galbas, Straky et al. 2024), often highly specific to the mouse model (Zacchigna, Paldino et al. 2021 (Rottman, Ni et al. 2007) and typically derived from small sample sizes (Stypmann, Engelen et al. 2006) (Vinhas, Araújo et al. 2013).

In this multicenter study, we have established reference ranges using an exceptionally large TTE dataset comprising more than 15,000 wildtype control mice from the International Mouse Phenotyping Consortium (IMPC). Rather than being highly specific to a single mouse model, our approach incorporates C57/BL6N substrains, enabling broader applicability across multiple related substrains. The goal of the IMPC is to extend the functional annotation of the mammalian genome via the large-scale production and phenotypic characterization of single gene knockout mouse strains for all protein coding genes. The phenotypic pipeline used to characterize these knockout strains included cardiac morphological and functional assessment using TTE. For each knockout strain characterized, age, sex and genetic background matched wildtype control animals were also assessed. The TTE data from these C57BL/6N wildtype control mice hold extraordinary value and represent the focus of the current study.

Thus, this study represents a large mouse data set and allows the crucial understanding of the effects of sex, age, and anesthesia on echocardiograms in mice. To this end, we introduced a stepwise refinement of the data analysis and started with an in-depth assessment of the variability of 12 TTE parameters gathered in the IMPC. We identified nine clinically relevant TTE parameters that were highly robust and had low variability. Mainly for methodological reasons, the respiration rate with high variability was excluded here, but LVAWd and LVAWs are the exceptions. These parameters were only collected by a minority of centers and therefore not included in the overall evaluation, but the values are made available in full in the supplemental materials.

By the nature of the multi-centre, large-scale IMPC approach, there are several confounding factors influencing the data. Understanding the sex, age, body weight, and anesthesia-related impact on TTE is crucial for ensuring robust reference values. In this study, we were able to show by a relative importance analysis that anesthetics are the main predictor with the largest proportion of explained variance in the model. Consequently, the entire data must be split according to anesthetic regime, as implemented here for conscious, anesthetized using isoflurane, or anesthetized using tribromoethanol states. Even with sex, body weight and age contributing little to the variance, understanding the impact of these factors is crucial for ensuring robust reference ranges.

Evidence indicates that sex-related factors interact in generating differences in cardiovascular outcomes in human (Regitz-Zagrosek and Gebhard 2023) and mice (Karp, Mason et al. 2017). In the current study, male and female mice generally presented with comparable reference ranges with negligible sexual dimorphism.

There may, however, be small sex differences for some parameters depending on the anesthetic agent. This observation is of key importance, and in part consistent with previous mouse data (Karp, Mason et al. 2017). Whilst sexual dimorphism was not overtly apparent in inbred mice in the absence of any environmental, pharmacological or genetic perturbations, the literature clearly supports sex differences in cardiovascular health, disease and risk prevention (Mensah and Fuster 2022, Regitz-Zagrosek and Gebhard 2023). Hence, our recommendation is that both sexes are included in any experimental design assuming that post-treatment we may detect sex differences.

Volatile anesthetics induce a dose-dependent decrease in myocardial contractility and cardiac loading conditions in mice and humans (Pagel 2013, Qin and Zhou 2023). These anesthetic effects reduce the metabolic activity and oxygen consumption of the heart. These protective properties can have a beneficial role on the myocardial oxygen balance during clinical anesthesia, surgery and post-operatively (De Hert 2006, Lin and Symons 2010, Abraham, Elliott et al. 2023). Our observations are that presence of anesthesia matters. We confirm a decreased heart rate in anesthetized mice and go on to reveal distinctions in isoflurane inhalation anesthesia and intraperitoneal injected tribromoethanol induced anesthesia. These distinctions are of central importance and impressively captured here. The known reduction in myocardial contractility under anesthesia is mirrored by increased left ventricular end-diastolic and end-systolic diameter and consequently decreased stroke volume, ejection fraction and fractional shortening, independent of age, anesthetic regime and sex of the mice. Following the calculation of cardiac output (CO = SV x HR), negligible anesthesia-related effects were anticipated, as LA mice exhibit a decrease in heart rate (HR) that is compensated by an increase in stroke volume (SV), resulting in comparable CO values to reference ranges. To emphasize the anesthesia distinctions, we mapped the effects of three different states (conscious, isoflurane and tribromoethanol anesthesia) on nine TTE parameters in detail and present anesthesia-specific reference values.

Cardiac aging is a complex and naturally inescapable process denominated by various molecular hallmarks leading to left ventricular hypertrophy, fibrosis, and end-diastolic dysfunction (Abdellatif, Rainer et al. 2023), where the mouse closely recapitulates the human situation (Dai and Rabinovitch 2009). Herein, we did observe a decrease in HR, an important determinant of sinoatrial node dysfunction with age (Peters, Sharpe et al. 2020), in 63-week-old mice (LA) compared to 12-week-old (EA) mice. Of interest, we identified age-related effects on end-systolic function and left ventricular (LV) size. In particular, we observed enhancement in end-systolic and end-diastolic LV dimensions with preserved LV posterior wall thickness and a reduction in the LV capacity demonstrated by decreased ejection fraction and fractional shortening with increased stroke volume and cardiac output in conscious and isoflurane-anesthetized LA mice of both sexes. These age-related cardiac characteristics were, however, marginal, and rather attributable to disease-free cardiovascular aging with no signs of pathophysiological cardiac remodeling. An age-related decline in myocardial elasticity through collagen fiber remodeling in combination with a drop in myocardial contractility caused by the diminution of myocardial cells and lower efficiency calcium channels provide rational explanations for the TTE parameter shifts observed herein in agreement with previous mouse studies (Xie, Fuchs et al. 2022). To this end, we present essential reference ranges that are critical for the assessment of normal age-related but disease-free cardiovascular ageing in mice (Obas and Vasan 2018) (Tracy, Rowe et al. 2020). Extending the aging period beyond 63 weeks, combined with interventions, could impact TTE parameters, though this was not part of the present study.

In the IMPC, we control for genetic diversity by using C57BL/6N inbred background substrains thereby focusing our comparison on the genetic perturbation of interest i.e. the single gene that is knocked out on this common genetic background. The transferability from the C57BL/6N background used here, however, was demonstrated by independently validating the ranges using data from a broad spectrum of non-IMPC C57BL/6N and C57BL/6J mice, and other inbred and wild-derived inbred strains ((Threadgill, Miller et al. 2011, Zacchigna, Paldino et al. 2021) https://phenome.jax.org/projects/Eumorphia6; https://phenome.jax.org/projects/Jaxwest1). This validation indicates that C57BL/6N-based reference values represent a robust and comprehensive indicator of normality for many strains and can be used as a starting point for experimental investigations of cardiac function in the mouse. A subset of outlier strain-parameter combinations was identified, for example, LVID;s, LVID;d, EF and FS in PWK/PhJ mice fell below the reported C57BL/6N-based reference range. The particularly small body weight of this wild-derived genetically diverse strain (Threadgill, Miller et al. 2011) (Kollmus, Fuchs et al. 2020) is consistent with a smaller but healthy heart and therefore explains the decreased LV inner dimensions.

This independent validation is also valid for reference values obtained in the ESC study using control/healthy C57BL/6 mice. A subset of outliers of the herein C57BL/6N-based reference values were identified, for example, LVIDs and LVIDd fell below and, as a consequence, EF and FS fell above the ESC reference values. Underlying data differed in key points such as data volume (300 versus 8.000 mice), anesthesia (isoflurane or sevoflurane versus isoflurane anesthesia), and age range (8-weeks to 3 months versus 12-weeks) potential factors that may explain the discrepancies between some parameter-specific reference values.

Each study comes with limitations. Echocardiography is performed at 2D M-mode, parasternal short-axis view only. Other echocardiographic views were omitted herein, pulse-wave Doppler and tissue Doppler were not in the scope of the IMPC large-scale high throughput phenotyping study. End-diastolic and end-systolic anterior wall thickness (LVAWd, LVAWs) was not entirely recorded at all contributing centers, yet the data is provided in full in Supplemental Table 3, stratified for sex and age. Body temperature was measured in all mice with anesthesia using a rectal probe and maintained within its physiological range (36°C–38°C) using a dedicated heating pad. Supplemental Figure 1 presents partial body temperature data to showcase the overall echocardiography conditions under anesthesia. The limiting factors for the tribromoethanol reference range data are that it was generated for 12-week-old mice only and the group size was the smallest of all conditions reported herein [450 mice distributed equally between sex (n = 227 female; n = 223 male)]. Solberg and colleagues (Solberg 1987), however, report that for a reliable estimate, a minimum of 120 values should be included for any reference range calculation. The sample size we used for tribromoethanol far exceeds this minimum and should therefore yield a representative range. The reference ranges are limited to the techniques and anesthetics described and are not intended for other imaging methodologies, such as MRI, or other anesthetic agents, such as ketamine.

Lack of transferability from mouse models to patients is the absence of standards and minimum requirements to ensure reliable and accurate assessment of cardiac function in mice. The reference ranges reported herein can be used to demarcate typical values for an experimental control group of mice on a C57BL/6N genetic background, for a given sex and age. They are not a substitute for contemporaneous control groups in any experimental design, but they indicate the likely values of that control group, thereby acting as a quality assurance tool. These reference ranges provide the information necessary to assess the changes in TTE parameters resulting from pharmacological, environmental, or genetic perturbations for experiments conducted on the commonly used C57BL/6N genetic background.

## Author contributions

Experiments, Data QC and Analysis: Marina Kan, Federico López Gómez, Robert Wilson, Susan Marschall, Mayra Monroy, Sheraz Pasha, Christopher Ward, Clare Norris, Dale Pimm, Sara Wells, Michelle Stewart, Eva Zábrodská, David Pajuelo Reguera, Zuzana Nichtova, Jan Prochazka, Hamid Meziane, Ghina Bou About, Mohammed Selloum,

Bioinformatics: Manuela A Oestereicher, and Schneltzer Elida and Helen Parkinson

Manuscript Design & Preparation: Manuela A Oestereicher; Christopher S. Ward and Nadine Spielmann

Study Design & Supervision Manuela A Oestereicher, Nadine Spielmann, Sara Wells, Michelle Stewart, Jan Prochazka, Radislav Sedláček, Yann Herault, Jason Heaney, Helmut Fuchs, Valerie Gailus-Durner; Martin Hrabe de Angelis, Xiang Gao, Christopher Ward,

PI on grant: Jason Heaney, Sara Wells, Radislav Sedláček, Yann Herault, Xiang Gao, Martin Hrabe de Angelis

Review the manuscript: Christopher S. Ward; Helmut Fuchs, Valerie Gailus-Durner; Jason Heaney, Martin Hrabe de Angelis, Yann Herault, Radislav Sedláček, Elida Schnelzer, Helen Parkinson

Mouse Production:

Phenotyping & Data Acquisition:

Data QC and Analysis:

Bioinformatic: Manuela A Oestereicher, Schneltzer Elida and Helen Parkinson

Manuscript Design & Preparation: Manuela A Oestereicher; Christopher S. Ward and Nadine Spielmann

Design & Study Supervision: Manuela A Oestereicher and Nadine Spielmann Review the manuscript: Christopher S. Ward; Helmut Fuchs, Valerie Gailus-Durner;

## Acknowledgements

Hands-on mouse IMPC phenotyping has been carried out by a large number of laboratory staff with superb experimental skills and unsurpassed dedication. The authors thank these individuals for their contribution to the IMPC data used herein.

## REFERENCES

1. (2008). Coefficient of Variation. The Concise Encyclopedia of Statistics. New York, NY, Springer New York: 95–96.

2. (2014). “A meta-analysis of echocardiographic measurements of the left heart for the development of normative reference ranges in a large international cohort: the EchoNoRMAL study.” Eur Heart J Cardiovasc Imaging 15(3): 341–348. AIM: To develop age-, sex-, and ethnic-appropriate normative reference ranges for standard echocardiographic measurements of the left heart by combining echocardiographic measurements obtained from adult volunteers without clinical cardiovascular disease or significant cardiovascular risk factors, from multiple studies around the world. METHODS AND RESULTS: The Echocardiographic Normal Ranges Meta-Analysis of the Left heart (EchoNoRMAL) collaboration was established and population-based data sets of echocardiographic measurements combined to perform an individual person data meta-analysis. Data from 43 studies were received, representing 51 222 subjects, of which 22 404 adults aged 18-80 years were without clinical cardiovascular or renal disease, hypertension or diabetes. Quantile regression or an appropriate parametric regression method will be used to derive reference values at the 5th and 95th centile of each measurement against age. CONCLUSION: This unique data set represents a large, multi-ethnic cohort of subjects resident in a wide range of countries. The resultant reference ranges will have wide applicability for normative data based on age, sex, and ethnicity.

3. (2015). “Ethnic-Specific Normative Reference Values for Echocardiographic LA and LV Size, LV Mass, and Systolic Function: The EchoNoRMAL Study.” JACC Cardiovasc Imaging 8(6): 656–665. OBJECTIVES: This study sought to derive age-, sex-, and ethnic-appropriate adult reference values for left atrial (LA) and left ventricular (LV) dimensions and volumes, LV mass, fractional shortening, and ejection fraction (EF) derived from geographically diverse population studies. BACKGROUND: The current recommended reference values for measurements from echocardiography may not be suitable to the diverse world population to which they are now applied. METHODS: Population-based datasets of echocardiographic measurements from 22,404 adults without clinical cardiovascular or renal disease, hypertension, or diabetes were combined in an individual person data meta-analysis. Quantile regression was used to derive reference values at the 95th percentile (upper reference value [URV]) and fifth percentile (lower reference value [LRV]) of each measurement against age (treated as linear), separately within sex and ethnic groups. RESULTS: The URVs for left ventricular end-diastolic volume (LVEDV), LV end-systolic volume, and LV stroke volume (SV) were highest in Europeans and lowest in South Asians. Important sex and ethnic differences remained after indexation by body surface area or height for these measurements, as well as for the LRV for SV. LVEDV and SV decreased with increasing age for all groups. Importantly, the LRV for EF differed by ethnicity; there was a clear apparent difference between Europeans and Asians. The URVs for LV end-diastolic diameter and LV end-systolic diameter were higher for Europeans than those for East Asian, South Asian, and African people, particularly among men. Similarly, the URVs for LA diameter and volume were highest for Europeans. CONCLUSIONS: Sex– and/or ethnic-appropriate echocardiographic reference values are indicated for many measurements of LA and LV size, LV mass, and EF. Reference values for LV volumes and mass also differ across the age range.

4. Abdellatif, M., et al. (2023). “Hallmarks of cardiovascular ageing.” Nature Reviews Cardiology 20(11): 754–777. Normal circulatory function is a key determinant of disease-free life expectancy (healthspan). Indeed, pathologies affecting the cardiovascular system, which are growing in prevalence, are the leading cause of global morbidity, disability and mortality, whereas the maintenance of cardiovascular health is necessary to promote both organismal healthspan and lifespan. Therefore, cardiovascular ageing might precede or even underlie body-wide, age-related health deterioration. In this Review, we posit that eight molecular hallmarks are common denominators in cardiovascular ageing, namely disabled macroautophagy, loss of proteostasis, genomic instability (in particular, clonal haematopoiesis of indeterminate potential), epigenetic alterations, mitochondrial dysfunction, cell senescence, dysregulated neurohormonal signalling and inflammation. We also propose a hierarchical order that distinguishes primary (upstream) from antagonistic and integrative (downstream) hallmarks of cardiovascular ageing. Finally, we discuss how targeting each of the eight hallmarks might be therapeutically exploited to attenuate residual cardiovascular risk in older individuals.

5. Abraham, A. S., et al. (2023). “Intra-operative anesthetic induced myocardial protection during cardiothoracic surgery: a literature review.” J Thorac Dis 15(12): 7042–7049. BACKGROUND AND OBJECTIVE: Myocardial protection involves limiting the metabolic activity and oxygen consumption of the heart, thus enabling surgery to proceed with minimal blood loss while reducing the level of ischemic injury. It was this concept that allowed for the development of the open-heart surgical technique. We know myocardial ischemia and reperfusion injury are both detrimental, thus developing strategies to mitigate this can help reduce peri-operative morbidity and mortality. In this review, we will mainly be addressing the anesthetic considerations for myocardial protection, along with discussing potential future research which can help expand the field. METHODS: We searched the PubMed database for relevant studies dating from 2004-2022. In total, 18 studies were deemed suitable for this literature review. KEY CONTENT AND FINDINGS: Studies have demonstrated cardioprotective effects with use of the volatile agents and propofol, mainly with respect to lower levels of inflammatory markers such as creatine kinase (CK)-MB and troponin I (TnI)/troponin T (TnT). The data is lacking regarding protective effects of dexmedetomidine and lidocaine, hence we cannot recommend either agent at present. CONCLUSIONS: Myocardial protection with respect to the anesthetic agents have been extensively studied over the past two decades, some routinely used drugs such as the volatile agents, propofol and opiates have demonstrated a cardioprotective role. The ideal dosing regimen and duration are areas of research that can be studied further. The data for the other anesthetic adjuncts such as lidocaine, dexmedetomidine along with use of regional anesthesia is still equivocal. Alongside advances in anesthesia, we believe surgical research looking into optimal cardioplegia solutions will also help improve myocardial protection in the future.

6. Ahn, Y., et al. (2024). “Reference ranges of computed tomography-derived strains in four cardiac chambers.” PLoS One 19(6): e0303986. Research on cardiovascular diseases using CT-derived strain is gaining momentum, yet there is a paucity of information regarding reference standard values beyond echocardiography, particularly in cardiac chambers other than the left ventricle (LV). We aimed to compile CT-derived strain values from the four cardiac chambers in healthy adults and assess the impact of age and sex on myocardial strains. This study included 101 (mean age: 55.2 ± 9.0 years, 55.4% men) consecutive healthy individuals who underwent multiphase cardiac CT. CT-derived cardiac strains, including LV global and segmental longitudinal, circumferential, and transverse strains, left atrial (LA), right atrial (RA), and right ventricle (RV) strains were measured by the commercially available software. Strain values were classified and compared by their age and sex. The normal range of CT-derived LV global longitudinal strain (GLS), global circumferential strain (GCS), and global radial strain (GRS) were –20.2 ± 2.7%, –27.9 ± 4.1%, and 49.4 ± 12.1%, respectively. For LA, reservoir strain, pump strain, and conduit strain were 28.6 ± 8.5%, 13.2 ± 6.4%, and 15.5 ± 8.6%, respectively. The GLS of RA and RV were 27.9 ± 10.9% and –22.0 ± 5.7%, respectively. The absolute values of GLS of RA and RV of women were higher than that in men (32.4 ± 11.4 vs. 24.3 ± 9.1 and –25.2 ± 4.7 vs. –19.4 ± 5.0, respectively; p<0.001, both). Measurement of CT-derived strain in four cardiac chambers is feasible. The reference ranges of CT strains in four cardiac chambers can be used for future studies of various cardiac diseases using the cardiac strains.

7. Akoglu, H. (2018). “User’s guide to correlation coefficients.” Turk J Emerg Med 18(3): 91–93. When writing a manuscript, we often use words such as perfect, strong, good or weak to name the strength of the relationship between variables. However, it is unclear where a good relationship turns into a strong one. The same strength of r is named differently by several researchers. Therefore, there is an absolute necessity to explicitly report the strength and direction of r while reporting correlation coefficients in manuscripts. This article aims to familiarize medical readers with several different correlation coefficients reported in medical manuscripts, clarify confounding aspects and summarize the naming practices for the strength of correlation coefficients.

8. Boyd, A. C., et al. (2015). “Principles of transthoracic echocardiographic evaluation.” Nature Reviews Cardiology 12(7): 426–440. Transthoracic echocardiography utilizes both visual (qualitative) and quantitative tools to evaluate cardiac structure and functionTraditional techniques, such as two-dimensional and Doppler imaging, continue to provide the framework for the continual improvements in transthoracic echocardiographyTissue Doppler, strain, and torsion echocardiography, are new imaging techniques that provide accurate and reliable quantitative assessment of myocardial functionOther advanced techniques, including stress, contrast, and three-dimensional echocardiography, provide additional diagnostic and functional informationThe technological advances in transthoracic echocardiography are evolving at a rapid rate

9. Brown, S. D. M., et al. (2005). “EMPReSS: standardized phenotype screens for functional annotation of the mouse genome.” Nature genetics 37(11): 1155.

10. Cain, P. A., et al. (2009). “Age and gender specific normal values of left ventricular mass, volume and function for gradient echo magnetic resonance imaging: a cross sectional study.” BMC Med Imaging 9: 2. BACKGROUND: Knowledge about age-specific normal values for left ventricular mass (LVM), end-diastolic volume (EDV), end-systolic volume (ESV), stroke volume (SV) and ejection fraction (EF) by cardiac magnetic resonance imaging (CMR) is of importance to differentiate between health and disease and to assess the severity of disease. The aims of the study were to determine age and gender specific normal reference values and to explore the normal physiological variation of these parameters from adolescence to late adulthood, in a cross sectional study. METHODS: Gradient echo CMR was performed at 1.5 T in 96 healthy volunteers (11-81 years, 50 male). Gender-specific analysis of parameters was undertaken in both absolute values and adjusted for body surface area (BSA). RESULTS: Age and gender specific normal ranges for LV volumes, mass and function are presented from the second through the eighth decade of life. LVM, ESV and EDV rose during adolescence and declined in adulthood. SV and EF decreased with age. Compared to adult females, adult males had higher BSA-adjusted values of EDV (p = 0.006) and ESV (p < 0.001), similar SV (p = 0.51) and lower EF (p = 0.014). No gender differences were seen in the youngest, 11-15 year, age range. CONCLUSION: LV volumes, mass and function vary over a broad age range in healthy individuals. LV volumes and mass both rise in adolescence and decline with age. EF showed a rapid decline in adolescence compared to changes throughout adulthood. These findings demonstrate the need for age and gender specific normal ranges for clinical use.

11. Cerbu, M., et al. (2023). “M-Mode Echocardiography in Canine Veterinary Practice: A Comprehensive Review of Left Ventricular Measurements in 44 Different Dog Breeds.” Animals (Basel) 13(18). This review article focuses on the use of canine M-mode in veterinary medicine, specifically in assessing the left ventricle measurements in several breeds. It traces the historical development of echocardiography techniques, including A-mode, B-mode, and motion mode (M-mode), which provide accurate unidimensional records of cardiac structures. This article highlights the significance of M-mode measurements in diagnosing stage B2 of MMVD, where left ventricular end-diastolic internal diameter corrected with body weight (LVIDdN) is essential for identifying cardiac enlargement. It also explains the role of M-mode in diagnosing DCM, outlining criteria such as left ventricular dilatation. The authors emphasize the importance of breed-specific reference values for echocardiographic measurements due to variations in somatotype among dogs. This review provides a comprehensive table summarizing M-mode measurements of the left ventricle for 44 different dog breeds, including interventricular septum thickness, left ventricular internal diameter, and left ventricular posterior wall thickness during systole and diastole. This review’s methodology involves compiling data from various scientific literature sources, providing an extensive tabular representation of M-mode measurements for different breeds, ages, and sexes. Overall, this review highlights the critical role of M-mode echocardiography in diagnosing and managing cardiac diseases in dogs, underscores the importance of breed-specific reference values, and presents a comprehensive summary of M-mode measurements for various dog breeds, aiding both clinicians and researchers.

12. Dai, D.-F. and P. S. Rabinovitch (2009). “Cardiac Aging in Mice and Humans: The Role of Mitochondrial Oxidative Stress.” Trends in Cardiovascular Medicine 19(7): 213–220. Age is a major risk factor for cardiovascular diseases, not only because it prolongs exposure to several other cardiovascular risks, but also owing to intrinsic cardiac aging, which reduces cardiac functional reserve, predisposes the heart to stress, and contributes to increased cardiovascular mortality in the elderly. Intrinsic cardiac aging in the murine model closely recapitulates age-related cardiac changes in humans, including left ventricular hypertrophy, fibrosis, and diastolic dysfunction. Cardiac aging in mice is accompanied by accumulation of mitochondrial protein oxidation, increased mitochondrial DNA mutations, increased mitochondrial biogenesis, as well as decreased cardiac SERCA2 protein. All of these age-related changes are significantly attenuated in mice overexpressing catalase targeted to mitochondria. These findings demonstrate the critical role of mitochondrial reactive oxygen species in cardiac aging and support the potential application of mitochondrial antioxidants to cardiac aging and age-related cardiovascular diseases.

13. De Hert, S. G. (2006). “Volatile Anesthetics and Cardiac Function.” Seminars in Cardiothoracic and Vascular Anesthesia 10(1): 33–42. All volatile anesthetics have been shown to induce a dose-dependent decrease in myocardial contractility and cardiac loading conditions. These depressant effects decrease myocardial oxygen demand and may, therefore, have a beneficial role on the myocardial oxygen balance during myocardial ischemia. Recently, experimental evidence has clearly demonstrated that in addition to theseindirect protective effects, volatile anesthetic agents also havedirect protective properties against reversible and irreversible ischemic myocardial damage. These properties have not only been related to a direct preconditioning effect but also to an effect on the extent of reperfusion injury. The implementation of these properties during clinical anesthesia can provide an additional tool in the treatment or prevention, or both, of ischemic cardiac dysfunction in the perioperative period. In the clinical practice, these effects should be associated with improved cardiac function, finally resulting in a better outcome in patients with coronary artery disease. The potential application of these protective properties of volatile anesthetic agents in clinical practice is the subject of ongoing research. This review summarizes the current knowledge on this subject.

14. Dickinson, M. E., et al. (2016). “High-throughput discovery of novel developmental phenotypes.” Nature 537(7621): 508–514. Approximately one-third of all mammalian genes are essential for life. Phenotypes resulting from knockouts of these genes in mice have provided tremendous insight into gene function and congenital disorders. As part of the International Mouse Phenotyping Consortium effort to generate and phenotypically characterize 5,000 knockout mouse lines, here we identify 410 lethal genes during the production of the first 1,751 unique gene knockouts. Using a standardized phenotyping platform that incorporates high-resolution 3D imaging, we identify phenotypes at multiple time points for previously uncharacterized genes and additional phenotypes for genes with previously reported mutant phenotypes. Unexpectedly, our analysis reveals that incomplete penetrance and variable expressivity are common even on a defined genetic background. In addition, we show that human disease genes are enriched for essential genes, thus providing a dataset that facilitates the prioritization and validation of mutations identified in clinical sequencing efforts.

15. Dobrev, D. and X. H. T. Wehrens (2018). “Mouse Models of Cardiac Arrhythmias.” Circ Res 123(3): 332–334. Mouse models have been invaluable for delineating the contributions of specific genes and signaling pathways to the pathogenesis of cardiac arrhythmias. Considering that there are important differences between mice and humans, here we discuss the strengths and limitations of mouse models of cardiac arrhythmias.

16. Domínguez-Oliva, A., et al. (2023). “The Importance of Animal Models in Biomedical Research: Current Insights and Applications.” Animals (Basel) 13(7). Animal research is considered a key element in advance of biomedical science. Although its use is controversial and raises ethical challenges, the contribution of animal models in medicine is essential for understanding the physiopathology and novel treatment alternatives for several animal and human diseases. Current pandemics’ pathology, such as the 2019 Coronavirus disease, has been studied in primate, rodent, and porcine models to recognize infection routes and develop therapeutic protocols. Worldwide issues such as diabetes, obesity, neurological disorders, pain, rehabilitation medicine, and surgical techniques require studying the process in different animal species before testing them on humans. Due to their relevance, this article aims to discuss the importance of animal models in diverse lines of biomedical research by analyzing the contributions of the various species utilized in science over the past five years about key topics concerning human and animal health.

17. Douglas, P. S., et al. (2019). “2019 ACC/AHA/ASE Key Data Elements and Definitions for Transthoracic Echocardiography: A Report of the American College of Cardiology/American Heart Association Task Force on Clinical Data Standards (Writing Committee to Develop Clinical Data Standards for Transthoracic Echocardiography) and the American Society of Echocardiography.” Circulation: Cardiovascular Imaging 12(7): e000027.

18. Eriksen-Volnes, T., et al. (2023). “Normalized Echocardiographic Values From Guideline-Directed Dedicated Views for Cardiac Dimensions and Left Ventricular Function.” JACC Cardiovasc Imaging 16(12): 1501–1515. BACKGROUND: Continuous technologic development and updated recommendations for image acquisitions creates a need to update the current normal reference ranges for echocardiography. The best method of indexing cardiac volumes is unknown. OBJECTIVES: The authors used 2– and 3-dimensional echocardiographic data from a large cohort of healthy individuals to provide updated normal reference data for dimensions and volumes of the cardiac chambers as well as central Doppler measurements. METHODS: In the fourth wave of the HUNT (Trøndelag Health) study in Norway 2,462 individuals underwent comprehensive echocardiography. Of these, 1,412 (55.8% women) were classified as normal and formed the basis for updated normal reference ranges. Volumetric measures were indexed to body surface area and height in powers of 1 to 3. RESULTS: Normal reference data for echocardiographic dimensions, volumes, and Doppler measurements were presented according to sex and age. Left ventricular ejection fraction had lower normal limits of 50.8% for women and 49.6% for men. According to sex-specific age groups, the upper normal limits for left atrial end-systolic volume indexed to body surface area ranged from 44 mL/m(2) to 53 mL/m(2), and the corresponding upper normal limit for right ventricular basal dimension ranged from 43 mm to 53 mm. Indexing to height raised to the power of 3 accounted for more of the variation between sexes than indexing to body surface area. CONCLUSIONS: The authors present updated normal reference values for a wide range of echocardiographic measures of both left– and right-side ventricular and atrial size and function from a large healthy population with a wide age-span. The higher upper normal limits for left atrial volume and right ventricular dimension highlight the importance of updating reference ranges accordingly following refinement of echocardiographic methods.

19. Galbas, M. C., et al. (2024). “Cardiac dimensions and hemodynamics in healthy juvenile Landrace swine.” Cardiovasc Ultrasound 22(1): 3. BACKGROUND: Swine are frequently used as animal model for cardiovascular research, especially in terms of representativity of human anatomy and physiology. Reference values for the most common species used in research are important for planning and execution of animal testing. Transesophageal echocardiography is the gold standard for intraoperative imaging, but can be technically challenging in swine. Its predecessor, epicardial echocardiography (EE), is a simple and fast intraoperative imaging technique, which allows comprehensive and goal-directed assessment. However, there are few echocardiographic studies describing echocardiographic parameters in juvenile swine, none of them using EE. Therefore, in this study, we provide a comprehensive dataset on multiple geometric and functional echocardiographic parameters, as well as basic hemodynamic parameters in swine using EE. METHODS: The data collection was performed during animal testing in ten female swine (German Landrace, 104.4 ± 13.0 kg) before left ventricular assist device implantation. Hemodynamic data was recorded continuously, before and during EE. The herein described echocardiographic measurements were acquired according to a standardized protocol, encompassing apical, left ventricular short axis and long axis as well as epiaortic windows. In total, 50 echocardiographic parameters and 10 hemodynamic parameters were assessed. RESULTS: Epicardial echocardiography was successfully performed in all animals, with a median screening time of 14 min (interquartile range 11-18 min). Referring to left ventricular function, ejection fraction was 51.6 ± 5.9% and 51.2 ± 6.2% using the Teichholz and Simpson methods, respectively. Calculated ventricular mass was 301.1 ± 64.0 g, as the left ventricular end-systolic and end-diastolic diameters were 35.3 ± 2.5 mm and 48.2 ± 3.5 mm, respectively. The mean heart rate was 103 ± 28 bpm, mean arterial pressure was 101 ± 20 mmHg and mean flow at the common carotid artery was 627 ± 203 mL/min. CONCLUSION: Epicardial echocardiography allows comprehensive assessment of most common echocardiographic parameters. Compared to humans, there are important differences in swine with respect to ventricular mass, size and wall thickness, especially in the right heart. Most hemodynamic parameters were comparable between swine and humans. This data supports study planning, animal and device selection, reinforcing the three R principles in animal research.

20. Galderisi, M., et al. (2017). “Standardization of adult transthoracic echocardiography reporting in agreement with recent chamber quantification, diastolic function, and heart valve disease recommendations: an expert consensus document of the European Association of Cardiovascular Imaging.” European Heart Journal – Cardiovascular Imaging 18(12): 1301–1310. This European Association Cardiovascular Imaging (EACVI) Expert Consensus document aims at defining the main quantitative information on cardiac structure and function that needs to be included in standard echocardiographic report following recent ASE/EACVI chamber quantification, diastolic function, and heart valve disease recommendations. The document focuses on general reporting and specific pathological conditions such as heart failure, coronary artery and valvular heart disease, cardiomyopathies, and systemic diseases.Demographic data (age, body surface area, blood pressure, and heart rhythm and rate), type (vendor and model) of ultrasound system used and image quality need to be reported. In addition, measurements should be normalized for body size. Reference normal values, derived by ASE/EACVI recommendations, shall always be reported to differentiate normal from pathological conditions. This Expert Consensus document suggests avoiding the surveillance of specific variable using different ultrasound techniques (e.g. in echo labs with high expertise in left ventricular ejection fraction by 3D and not by 2D echocardiography). The report should be also tailored in relation with different cardiac pathologies, quality of images, and needs of the caregivers.The conclusion should be concise reflecting the status of left ventricular structure and function, the presence of left atrial and/or aortic dilation, right ventricular dysfunction, and pulmonary hypertension, leading to an objective communication with the patient health caregiver. Variation over time should be considered carefully, taking always into account the consistency of the parameters used for comparison.

21. Gao, S., et al. (2011). “Echocardiography in Mice.” Curr Protoc Mouse Biol 1: 71–83. Murine models have been utilized with increasing frequency mainly due to availability of genetically engineered models. With advancement in high spatial and temporal resolution, echocardiography is used extensively for the evaluation of cardiovascular function in murine models of cardiovascular disease. This review summarizes the general applications and methods involved in echocardiography used to study mouse models for cardiovascular research, based on 20 years of experience in our laboratory. The goal of this article is to provide a practical guide to the use of echo techniques in mice to evaluate cardiac systolic and diastolic function.

22. Grenacher, P. A. and C. C. Schwarzwald (2010). “Assessment of left ventricular size and function in horses using anatomical M-mode echocardiography.” J Vet Cardiol 12(2): 111–121. OBJECTIVE: To study the applicability of anatomical M-mode (AMM) for assessment of left ventricular (LV) size and function in horses, evaluate agreement with conventional M-mode (CMM), determine reliability, and establish reference intervals for AMM measurements. ANIMALS: 98 horses; 13.1 +/-5.6 years; 538 +/-78 kg. METHODS: Two-dimensional and M-mode recordings were analyzed retrospectively. Standard LV dimensions and indices of LV function, including time intervals, were measured in CMM and compared with AMM studies in long-axis (lx) and short-axis (sx) views. RESULTS: The percentages of measureable cycles were 99%, 97%, and 90% for routine LV studies in CMM(sx), AMM(sx), and AMM(lx) mode. For time intervals, >or= 93% of cycles could be measured using AMM compared to a maximum of 77% using CMM. AMM(sx) measurements agreed well with CMM(sx) measurements for LV studies; the agreement of AMM(lx) with CMM(sx) was markedly lower. The LV ejection time and the duration of electromechanical systole, but not the LV pre-ejection period and the index of myocardial performance, showed fair agreement between methods. Intraobserver and interobserver measurement variabilities were low for most variables. CONCLUSIONS: AMM can replace CMM for assessment of LV dimensions in horses, but is not recommended for measurement of time intervals.

23. Groemping, U. (2006). “Relative Importance for Linear Regression in R: The Package relaimpo.” Journal of Statistical Software 17(1): 1–27. Relative importance is a topic that has seen a lot of interest in recent years, particularly in applied work. The R package relaimpo implements six different metrics for assessing relative importance of regressors in the linear model, two of which are recommended – averaging over orderings of regressors and a newly proposed metric (Feldman 2005) called pmvd. Apart from delivering the metrics themselves, relaimpo also provides (exploratory) bootstrap confidence intervals. This paper offers a brief tutorial introduction to the package. The methods and relaimpo’s functionality are illustrated using the data set swiss that is generally available in R. The paper targets readers who have a basic understanding of multiple linear regression. For the background of more advanced aspects, references are provided.

24. Harkness, A., et al. (2020). “Normal Reference Intervals for Cardiac Dimensions and Function for Use in Echocardiographic Practice: A Guideline from the British Society of Echocardiography.” Echo Research & Practice 7(1): G1–G18. This guideline presents reference limits for use in echocardiographic practice, updating previous guidance from the British Society of Echocardiography. The rationale for change is discussed, in addition to how the reference intervals were defined and the current limitations to their use. The importance of interpretation of echocardiographic parameters within the clinical context is explored, as is grading of abnormality. Each of the following echo parameters are discussed and updated in turn: left ventricular linear dimensions and LV mass; left ventricular volumes; left ventricular ejection fraction; left atrial size; right heart parameters; aortic dimensions; and tissue Doppler imaging. There are several important conceptual changes to the assessment of the heart’s structure and function within this guideline. New terminology for left ventricular function and left atrial size are introduced. The British Society of Echocardiography has advocated a new approach to the assessment of the aortic root, the right heart, and clarified the optimal methodology for assessment of LA size. The British Society of Echocardiography has emphasized a preference to use, where feasible, indexed measures over absolute values for any chamber size.

25. Heidenreich, P. (2023). “What Is a Normal Left Ventricular Ejection Fraction?” Circulation 148(9): 750–752.

26. Jasaityte, R., et al. (2013). “Normal changes of LV contractility with ageing: an echocardiographic study.” European Heart Journal 34(suppl_1). Purpose: Data about the natural changes of intrinsic LV contractility with age are sparse. We have recently proposed a novel echocardiographic approach to estimate LV contractility non-invasively by the slope of segmental passive stretch (preS) and systolic strain (SS) relationship. In this study we apply it in healthy volunteers of various age to detect how LV contractility changes with age in healthy hearts.Methods: In 54 healthy subjects from 21 to 70 years of age TDI (FR ∼200 Hz) of 6 LV walls were acquired. For each wall, base mid and apical regional strain curves were extracted with custom software, setting the reference point at the onset of P wave on ECG. PreS was measured as the peak positive strain after the P wave and SS as total systolic shortening. For each subject a linear regression line was estimated through 18 segmental preS and SS values. The study population was divided in 5 age groups by decades of years. Mean preS-SS relations were determined as an average slope and intercept in each group.Results: No differences of LV ejection fraction, systolic blood pressure or heart rate were observed between the groups. Mitral inflow A wave velocity and E wave deceleration time increased and E/A decreased significantly with age. A significant increase of LV PreS was observed with age, whereas SS remained unchanged. PreS and SS correlated in all subjects (mean r2=0.68). The slopes of PreS-SS relationship did not differ between the age groups while the intercept increased decreased slightly, but not significantly with increasing age (Fig. 1).Figure 1. preS-SS relationships and meansConclusion: The slope of the stretch-strain relationship did not change with age showing that intrinsic LV contractility remains unaltered in a healthy aging heart. It is in concordance with the results of recently published invasive study.

27. Karp, N. A., et al. (2017). “Prevalence of sexual dimorphism in mammalian phenotypic traits.” Nature Communications 8(1): 15475. The role of sex in biomedical studies has often been overlooked, despite evidence of sexually dimorphic effects in some biological studies. Here, we used high-throughput phenotype data from 14,250 wildtype and 40,192 mutant mice (representing 2,186 knockout lines), analysed for up to 234 traits, and found a large proportion of mammalian traits both in wildtype and mutants are influenced by sex. This result has implications for interpreting disease phenotypes in animal models and humans.

28. Kollmus, H., et al. (2020). “A comprehensive and comparative phenotypic analysis of the collaborative founder strains identifies new and known phenotypes.” Mamm Genome 31(1-2): 30–48. The collaborative cross (CC) is a large panel of mouse-inbred lines derived from eight founder strains (NOD/ShiLtJ, NZO/HILtJ, A/J, C57BL/6J, 129S1/SvImJ, CAST/EiJ, PWK/PhJ, and WSB/EiJ). Here, we performed a comprehensive and comparative phenotyping screening to identify phenotypic differences and similarities between the eight founder strains. In total, more than 300 parameters including allergy, behavior, cardiovascular, clinical blood chemistry, dysmorphology, bone and cartilage, energy metabolism, eye and vision, immunology, lung function, neurology, nociception, and pathology were analyzed; in most traits from sixteen females and sixteen males. We identified over 270 parameters that were significantly different between strains. This study highlights the value of the founder and CC strains for phenotype-genotype associations of many genetic traits that are highly relevant to human diseases. All data described here are publicly available from the mouse phenome database for analyses and downloads.

29. Kondo, T., et al. (2023). “Clinical Characteristics and Outcomes in Patients With Heart Failure: Are There Thresholds and Inflection Points in Left Ventricular Ejection Fraction and Thresholds Justifying a Clinical Classification?” Circulation 148(9): 732–749. BACKGROUND: Recent guidelines proposed a classification for heart failure (HF) on the basis of left ventricular ejection fraction (LVEF), although it remains unclear whether the divisions chosen were biologically rational. Using patients spanning the full range of LVEF, we examined whether there was evidence of LVEF thresholds in patient characteristics or inflection points in clinical outcomes. METHODS: Using patient-level information, we created a merged dataset of 33 699 participants who had been enrolled in 6 randomized controlled HF trials including patients with reduced and preserved ejection fraction. The relationship between the incidence of all-cause death (and specific causes of death) and HF hospitalization, and LVEF, was evaluated using Poisson regression models. RESULTS: As LVEF increased, age, the proportion of women, body mass index, systolic blood pressure, and prevalence of atrial fibrillation and diabetes increased, whereas ischemic pathogenesis, estimated glomerular filtration rate, and NT-proBNP (N-terminal pro-B-type natriuretic peptide) decreased. As LVEF increased >50%, age and the proportion of women continued to increase, and ischemic pathogenesis and NT-proBNP decreased, but other characteristics did not change meaningfully. The incidence of most clinical outcomes (except noncardiovascular death) decreased as LVEF increased, with a LVEF inflection point of around 50% for all-cause death and cardiovascular death, around 40% for pump failure death, and around 35% for HF hospitalization. Higher than those thresholds, there was little further decline in the incidence rate. There was no evidence of a J-shaped relationship between LVEF and death; no evidence of worse outcomes in patients with high-normal (“supranormal”) LVEF. Similarly, in a subset of patients with echocardiographic data, there were no structural differences in patients with a high-normal LVEF suggestive of amyloidosis, and NT-proBNP levels were consistent with this conclusion. CONCLUSIONS: In patients with HF, there was a LVEF threshold of around 40% to 50% where the pattern of patient characteristics changed, and event rates began to increase compared with higher LVEF values. Our findings provide evidence to support current upper LVEF thresholds defining HF with mildly reduced ejection fraction on the basis of prognosis. REGISTRATION: URL: https://www.CLINICALTRIALS:gov; Unique identifiers: NCT00634309, NCT00634400, NCT00634712, NCT00095238, NCT01035255, NCT00094302, NCT00853658, and NCT01920711.

30. Konrad, D., et al. (2000). “Echocardiography, color-coded Doppler imaging, and abdominal sonography, a non-invasive method for investigation of heart and aortic morphology and function in female gottingen minipigs: method and reference values for M-mode, B-mode, and flow parameters.” Comp Med 50(4): 405–409. OBJECTIVE: The aim of the study reported here was to set up a method for echocardiography (EC) and abdominal sonography and to obtain EC reference values for left ventricular (LV) morphology and function and sonographic abdominal aortic morphology, function, and flow values in conscious, unsedated Gottingen minipigs. METHODS: Applying a standardized investigation procedure, the following parameters were measured by use of M-mode EC, color-coded Doppler imaging, and B-mode sonography, or were calculated, in 58 female minipigs: LV end-diastolic and end-systolic diameter, interventricular septum thickness, LV caudal wall thickness, LV end-diastolic volume and end-systolic volume, fractional shortening, ejection fraction, and percentage of thickening of interventricular septum and LV caudal wall. In addition, morphology, pulsatility, flow values, and flow patterns in the abdominal aorta were recorded or calculated during abdominal sonography and color-coded Doppler imaging. RESULTS: Variable EC values were obtained due to individual variations of motor activity. Variation could be reduced by accustoming the animals to a standardized investigation procedure. Reference values could be obtained for EC, partially indicating clear correlation with body weight. Color-coded Doppler and Doppler spectra did not indicate flow disturbances in large arterial abdominal vessels. CONCLUSIONS: Results indicate that handling during EC and sonography can cause discomfort in unsedated minipigs that may interfere with recording of valid reference values for functional cardiac parameters in young animals. Accustoming the animals to a standardized investigation procedure reduces stress to a satisfactory level and enables data recording. Thus the minipig is considered suitable for assessment of cardiovascular parameters in experimental or toxicologic studies.

31. Lancellotti, P., et al. (2013). “Normal Reference Ranges for Echocardiography: rationale, study design, and methodology (NORRE Study).” Eur Heart J Cardiovasc Imaging 14(4): 303–308. BACKGROUND: Availability of normative reference values for cardiac chamber dimensions, volumes, mass, and function is a prerequisite for the accurate application of echocardiography for both clinical and research purposes. However, due to the lack of consistency in current echocardiographic ‘reference values’, their use for clinical decision-making remains questionable. AIMS: The aim of the ‘Normal Reference Ranges for Echocardiography Study (NORRE Study)’ is to obtain a set of ‘normal values’ for cardiac chamber geometry and function in a large cohort of healthy Caucasian individuals aged over a wide range of ages (25-75 years) using both conventional and advanced echocardiographic techniques. METHODS: The NORRE Study is a large prospective, observational multicentre study in which transthoracic echocardiographic studies will be acquired in 22 laboratories accredited by the European Association of Cardiovascular Imaging and in one laboratory in the USA accredited by ICAEL. The final sample size has been estimated in 1100 normal subjects in whom M-mode, 2D, and 3D imaging, colour Doppler, pulsed-wave Doppler, pulsed-wave tissue Doppler, and colour tissue Doppler imaging data will be obtained. All studies will be sent to a central echocardiographic core laboratory for quantitative analysis. Multiple studies will be performed for reproducibility analysis. CONCLUSION: After completion of the NORRE Study, uniform reference limits according to age, gender, and anthropometric parameters will be available to standardize the quantitative interpretation of echocardiography.

32. Lin, E. and J. A. Symons (2010). “Volatile anaesthetic myocardial protection: a review of the current literature.” HSR Proc Intensive Care Cardiovasc Anesth 2(2): 105–109. Ischaemic preconditioning is a powerful innate adaptive phenomenon whereby brief periods of sublethal ischaemia result in marked tolerance to subsequent lethal ischaemia. Halogenated anaesthetics have been shown to mimic ischaemic preconditioning, modifying and attenuating ischaemia reperfusion injury. This review aims to present the current animal and human data, discuss the possible mechanisms of action and review the clinical evidence for volatile anaesthetic-induced myocardial protection. There is class Ia evidence for the myocardial protective properties of sevoflurane and desflurane in low risk patients undergoing coronary artery bypass grafting surgery. These volatile anaesthetics have been shown to improve clinical outcomes and health economics following cardiac surgery, reducing intensive care and hospital stay. The evidence for the benefit of volatile anaesthetics in non-cardiac surgery is less robust and further large randomized controlled trials are required to elucidate this question.

33. Lindsey, M. L., et al. (2021). “Guidelines for in vivo mouse models of myocardial infarction.” Am J Physiol Heart Circ Physiol 321(6): H1056–h1073. Despite significant improvements in reperfusion strategies, acute coronary syndromes all too often culminate in a myocardial infarction (MI). The consequent MI can, in turn, lead to remodeling of the left ventricle (LV), the development of LV dysfunction, and ultimately progression to heart failure (HF). Accordingly, an improved understanding of the underlying mechanisms of MI remodeling and progression to HF is necessary. One common approach to examine MI pathology is with murine models that recapitulate components of the clinical context of acute coronary syndrome and subsequent MI. We evaluated the different approaches used to produce MI in mouse models and identified opportunities to consolidate methods, recognizing that reperfused and nonreperfused MI yield different responses. The overall goal in compiling this consensus statement is to unify best practices regarding mouse MI models to improve interpretation and allow comparative examination across studies and laboratories. These guidelines will help to establish rigor and reproducibility and provide increased potential for clinical translation.

34. Mensah, G. A. and V. Fuster (2022). “Sex and Gender Differences in Cardiovascular Health.” Journal of the American College of Cardiology 79(14): 1385–1387.

35. Muñoz-Fuentes, V., et al. (2018). “The International Mouse Phenotyping Consortium (IMPC): a functional catalogue of the mammalian genome that informs conservation.” Conserv Genet 19(4): 995–1005. The International Mouse Phenotyping Consortium (IMPC) is building a catalogue of mammalian gene function by producing and phenotyping a knockout mouse line for every protein-coding gene. To date, the IMPC has generated and characterised 5186 mutant lines. One-third of the lines have been found to be non-viable and over 300 new mouse models of human disease have been identified thus far. While current bioinformatics efforts are focused on translating results to better understand human disease processes, IMPC data also aids understanding genetic function and processes in other species. Here we show, using gorilla genomic data, how genes essential to development in mice can be used to help assess the potentially deleterious impact of gene variants in other species. This type of analyses could be used to select optimal breeders in endangered species to maintain or increase fitness and avoid variants associated to impaired-health phenotypes or loss-of-function mutations in genes of critical importance. We also show, using selected examples from various mammal species, how IMPC data can aid in the identification of candidate genes for studying a condition of interest, deliver information about the mechanisms involved, or support predictions for the function of genes that may play a role in adaptation. With genotyping costs decreasing and the continued improvements of bioinformatics tools, the analyses we demonstrate can be routinely applied.

36. Nyberg, J., et al. (2023). “Echocardiographic Reference Ranges of Global Longitudinal Strain for All Cardiac Chambers Using Guideline-Directed Dedicated Views.” JACC: Cardiovascular Imaging 16(12): 1516–1531.

37. Nyberg, J., et al. (2023). “Echocardiographic Reference Ranges of Global Longitudinal Strain for All Cardiac Chambers Using Guideline-Directed Dedicated Views.” JACC Cardiovasc Imaging 16(12): 1516–1531. BACKGROUND: Myocardial deformation by echocardiographic strain imaging is a key measurement in cardiology, providing valuable diagnostic and prognostic information. Reference ranges for strain should be established from large healthy populations with minimal methodologic biases and variability. OBJECTIVES: The aim of this study was to establish echocardiographic reference ranges, including lower normal limits of global strains for all 4 cardiac chambers, by guideline-directed dedicated views from a large healthy population and to evaluate the influence of subject-specific characteristics on strain. METHODS: In total, 1,329 healthy participants from HUNT4Echo, the echocardiographic substudy of the 4th wave of the Trøndelag Health Study, were included. Echocardiographic recordings specific for each chamber were optimized according to current recommendations. Two experienced sonographers recorded all echocardiograms using GE HealthCare Vivid E95 scanners. Analyses were performed by experts using GE HealthCare EchoPAC. RESULTS: The reference ranges for left ventricular (LV) global longitudinal strain and right ventricular free-wall strain were –24% to –16% and –35% to –17%, respectively. Correspondingly, left atrial (LA) and right atrial (RA) reservoir strains were 17% to 49% and 17% to 59%. All strains showed lower absolute values with higher age, except for LA and RA contractile strains, which were higher. The feasibility for strain was overall good (LV 96%, right ventricular 83%, LA 94%, and RA 87%). All chamber-specific strains were associated with age, and LV strain was associated with sex. CONCLUSIONS: Reference ranges of strain for all cardiac chambers were established based on guideline-directed chamber-specific recordings. Age and sex were the most important factors influencing reference ranges and should be considered when using strain echocardiography.

38. O’Riordan, C. E., et al. (2023). “Standardisation and future of preclinical echocardiography.” Mammalian Genome: 1–33.

39. Obas, V. and R. S. Vasan (2018). “The aging heart.” Clin Sci (Lond) 132(13): 1367–1382. As the elderly segment of the world population increases, it is critical to understand the changes in cardiac structure and function during the normal aging process. In this review, we outline the key molecular pathways and cellular processes that underlie the phenotypic changes in the heart and vasculature that accompany aging. Reduced autophagy, increased mitochondrial oxidative stress, telomere attrition, altered signaling in insulin-like growth factor, growth differentiation factor 11, and 5’-AMP-activated protein kinase pathways are among the key molecular mechanisms underlying cardiac aging. Aging promotes structural and functional changes in the atria, ventricles, valves, myocardium, pericardium, the cardiac conduction system, and the vasculature. We highlight the factors known to accelerate and attenuate the intrinsic aging of the heart and vessels in addition to potential preventive and therapeutic avenues. A greater understanding of the processes involved in cardiac aging may facilitate our ability to mitigate the escalating burden of CVD in older individuals and promote healthy cardiac aging.

40. Pagel, P. S. (2013). “Myocardial Protection by Volatile Anesthetics in Patients Undergoing Cardiac Surgery: A Critical Review of the Laboratory and Clinical Evidence.” Journal of Cardiothoracic and Vascular Anesthesia 27(5): 972–982.

41. Pearson, K. (1895). “Note on Regression and Inheritance in the Case of Two Parents.” Proceedings of the Royal Society of London 58: 240–242.

42. Peters, C. H., et al. (2020). “Cardiac Pacemaker Activity and Aging.” Annu Rev Physiol 82: 21–43. A progressive decline in maximum heart rate (mHR) is a fundamental aspect of aging in humans and other mammals. This decrease in mHR is independent of gender, fitness, and lifestyle, affecting in equal measure women and men, athletes and couch potatoes, spinach eaters and fast food enthusiasts. Importantly, the decline in mHR is the major determinant of the age-dependent decline in aerobic capacity that ultimately limits functional independence for many older individuals. The gradual reduction in mHR with age reflects a slowing of the intrinsic pacemaker activity of the sinoatrial node of the heart, which results from electrical remodeling of individual pacemaker cells along with structural remodeling and a blunted β-adrenergic response. In this review, we summarize current evidence about the tissue, cellular, and molecular mechanisms that underlie the reduction in pacemaker activity with age and highlight key areas for future work.

43. Pfaffenberger, S., et al. (2013). “Size Matters! Impact of Age, Sex, Height, and Weight on the Normal Heart Size.” Circulation: Cardiovascular Imaging 6(6): 1073–1079.

44. Qin, H. and J. Zhou (2023). “Myocardial Protection by Desflurane: From Basic Mechanisms to Clinical Applications.” J Cardiovasc Pharmacol 82(3): 169–179. Coronary heart disease is an affiction that is common and has an adverse effect on patients’ quality of life and survival while also raising the risk of intraoperative anesthesia. Mitochondria are the organelles most closely associated with the pathogenesis, development, and prognosis of coronary heart disease. Ion abnormalities, an acidic environment, the production of reactive oxygen species, and other changes during abnormal myocardial metabolism cause the opening of mitochondrial permeability transition pores, which disrupts electron transport, impairs mitochondrial function, and even causes cell death. Differences in reliability and cost-effectiveness between desflurane and other volatile anesthetics are minor, but desflurane has shown better myocardial protective benefits in the surgical management of patients with coronary artery disease. The results of myocardial protection by desflurane are briefly summarized in this review, and biological functions of the mitochondrial permeability transition pore, mitochondrial electron transport chain, reactive oxygen species, adenosine triphosphate-dependent potassium channels, G protein-coupled receptors, and protein kinase C are discussed in relation to the protective mechanism of desflurane. This article also discusses the effects of desflurane on patient hemodynamics, myocardial function, and postoperative parameters during coronary artery bypass grafting. Although there are limited and insufficient clinical investigations, they do highlight the possible advantages of desflurane and offer additional suggestions for patients.

45. Regitz-Zagrosek, V. and C. Gebhard (2023). “Gender medicine: effects of sex and gender on cardiovascular disease manifestation and outcomes.” Nature Reviews Cardiology 20(4): 236–247. Despite a growing body of evidence, the distinct contributions of biological sex and the sociocultural dimension of gender to the manifestations and outcomes of ischaemic heart disease and heart failure remain unknown. The intertwining of sex-based differences in genetic and hormonal mechanisms with the complex dimension of gender and its different components and determinants that result in different disease phenotypes in women and men needs to be elucidated. The relative contribution of purely biological factors, such as genes and hormones, to cardiovascular phenotypes and outcomes is not yet fully understood. Increasing awareness of the effects of gender has led to efforts to measure gender in retrospective and prospective clinical studies and the development of gender scores. However, the synergistic or opposing effects of sex and gender on cardiovascular traits and on ischaemic heart disease and heart failure mechanisms have not yet been systematically described. Furthermore, specific considerations of sex-related and gender-related factors in gender dysphoria or in heart–brain interactions and their association with cardiovascular disease are still lacking. In this Review, we summarize contemporary evidence on the distinct effects of sex and gender as well as of their interactions on cardiovascular disease and how they favourably or unfavourably influence the pathogenesis, clinical manifestations and treatment responses in patients with ischaemic heart disease or heart failure.

46. Robinson, N. B., et al. (2019). “The current state of animal models in research: A review.” International Journal of Surgery 72: 9–13. Animal models have provided invaluable information in the pursuit of medical knowledge and alleviation of human suffering. The foundations of our basic understanding of disease pathophysiology and human anatomy can largely be attributed to preclinical investigations using various animal models. Recently, however, the scientific community, citing concerns about animal welfare as well as the validity and applicability of outcomes, has called the use of animals in research into question. In this review, we seek to summarize the current state of the use of animal models in research.

47. Rottman, J. N., et al. (2007). “Echocardiographic evaluation of ventricular function in mice.” Echocardiography 24(1): 83–89. Ventricular dysfunction remains a hallmark of most cardiac disease. The mouse has become an essential model system for cardiovascular biology, and echocardiography an established tool in the study of normal and genetically altered mice. This review describes the measurement of ventricular function, most often left ventricular function, by echocardiographic methods in mice. Technical limitations related to the small size and rapid heart rate in the mouse initially argued for the performance of echocardiography under anesthesia. More recently, higher frame rates and smaller probes operating at higher frequencies have facilitated imaging of conscious mice in some, but not all, experimental protocols and conditions. Ventricular function may be qualitatively and quantitatively evaluated under both conditions. Particular detail is provided for measurement under conscious conditions, and measurement under conscious and sedated or anesthestized conditions are contrasted. Normal values for echocardiographic indices for the common C57BL/6 strain are provided. Diastolic dysfunction is a critical pathophysiologic component of many disease states, and progress in the echocardiographic evaluation of diastolic function is discussed. Finally, echocardiography exists among several competing imaging technologies, and these alternatives are compared.

48. Schober, P., et al. (2018). “Correlation Coefficients: Appropriate Use and Interpretation.” Anesth Analg 126(5): 1763–1768. Correlation in the broadest sense is a measure of an association between variables. In correlated data, the change in the magnitude of 1 variable is associated with a change in the magnitude of another variable, either in the same (positive correlation) or in the opposite (negative correlation) direction. Most often, the term correlation is used in the context of a linear relationship between 2 continuous variables and expressed as Pearson product-moment correlation. The Pearson correlation coefficient is typically used for jointly normally distributed data (data that follow a bivariate normal distribution). For nonnormally distributed continuous data, for ordinal data, or for data with relevant outliers, a Spearman rank correlation can be used as a measure of a monotonic association. Both correlation coefficients are scaled such that they range from –1 to +1, where 0 indicates that there is no linear or monotonic association, and the relationship gets stronger and ultimately approaches a straight line (Pearson correlation) or a constantly increasing or decreasing curve (Spearman correlation) as the coefficient approaches an absolute value of 1. Hypothesis tests and confidence intervals can be used to address the statistical significance of the results and to estimate the strength of the relationship in the population from which the data were sampled. The aim of this tutorial is to guide researchers and clinicians in the appropriate use and interpretation of correlation coefficients.

49. Solberg, H. E. (1987). “Approved recommendation (1986) on the theory of reference values. Part 1. The concept of reference values.” Clinica Chimica Acta 165(1): 111–118.

50. Stypmann, J., et al. (2006). “Age and gender related reference values for transthoracic Doppler-echocardiography in the anesthetized CD1 mouse.” Int J Cardiovasc Imaging 22(3-4): 353–362. OBJECTIVE: Doppler-echocardiography of the mouse has evolved to a commonly used technique in the past years as recent advances in imaging quality have substantially improved spatial and temporal resolution allowing the adaptation of this technique to murine models. Although mouse echocardiography is widely used, there is only little information on reference data for wild-type animals available, particularly in older mice. METHODS: We therefore established a database with echocardiographic reference-values in a large set of young (8 weeks) and older adult (52 weeks) Swiss type CD1-mice of either sex. We performed a complete Doppler-echocardiographic examination under light Ketamine-Xylazine-anesthesia. LV-mass was calculated and compared with necropsy heart weights to validate the LV-mass calculation. RESULTS: Doppler-echocardiographic measurements in mice were feasible to assess cardiac morphology and function. Sonomorphological and functional parameters hardly changed between the age of 12 and 52 weeks. Wall thickness, LV-mass and cardiac output were stable with aging. There was a good relative correlation between echocardiographically estimated LV-mass and necropsy heart weight although absolute values differed. There were no significant echocardiographic differences between male and female mice. CONCLUSIONS: The reference values established in this study can be useful in recording and quantifying pathological changes in murine models of cardiovascular diseases. There is hardly any change of cardiac function between the age of 12 and 52 weeks.

51. Team, R. C. (2022). R: A Language and Environment for Statistical Computing. R Foundation for Statistical Computing. https://www.R-project.org. **R Core Team (**2022).

52. Threadgill, D. W., et al. (2011). “The Collaborative Cross: A Recombinant Inbred Mouse Population for the Systems Genetic Era.” ILAR Journal 52(1): 24–31. The mouse is the most extensively used mammalian model for biomedical and aging research, and an extensive catalogue of laboratory resources is available to support research using mice: classical inbred lines, genetically modified mice (knockouts, transgenics, and humanized mice), selectively bred lines, consomics, congenics, recombinant inbred panels, outbred and heterogeneous stocks, and an expanding set of wild-derived strains. However, these resources were not designed or intended to model the heterogeneous human population or for a systematic analysis of phenotypic effects due to random combinations of uniformly distributed natural variants. The Collaborative Cross (CC) is a large panel of recently established multiparental recombinant inbred mouse lines specifically designed to overcome the limitations of existing mouse genetic resources for analysis of phenotypes caused by combinatorial allele effects. The CC models the complexity of the human genome and supports analyses of common human diseases with complex etiologies originating through interactions between allele combinations and the environment. The CC is the only mammalian resource that has high and uniform genomewide genetic variation effectively randomized across a large, heterogeneous, and infinitely reproducible population. The CC supports data integration across environmental and biological perturbations and across space (different labs) and time.

53. Threadgill, D. W., et al. (2011). “The collaborative cross: a recombinant inbred mouse population for the systems genetic era.” Ilar j 52(1): 24–31. The mouse is the most extensively used mammalian model for biomedical and aging research, and an extensive catalogue of laboratory resources is available to support research using mice: classical inbred lines, genetically modified mice (knockouts, transgenics, and humanized mice), selectively bred lines, consomics, congenics, recombinant inbred panels, outbred and heterogeneous stocks, and an expanding set of wild-derived strains. However, these resources were not designed or intended to model the heterogeneous human population or for a systematic analysis of phenotypic effects due to random combinations of uniformly distributed natural variants. The Collaborative Cross (CC) is a large panel of recently established multiparental recombinant inbred mouse lines specifically designed to overcome the limitations of existing mouse genetic resources for analysis of phenotypes caused by combinatorial allele effects. The CC models the complexity of the human genome and supports analyses of common human diseases with complex etiologies originating through interactions between allele combinations and the environment. The CC is the only mammalian resource that has high and uniform genomewide genetic variation effectively randomized across a large, heterogeneous, and infinitely reproducible population. The CC supports data integration across environmental and biological perturbations and across space (different labs) and time.

54. Tracy, E., et al. (2020). “Cardiac tissue remodeling in healthy aging: the road to pathology.” Am J Physiol Cell Physiol 319(1): C166–c182. This review aims to highlight the normal physiological remodeling that occurs in healthy aging hearts, including changes that occur in contractility, conduction, valve function, large and small coronary vessels, and the extracellular matrix. These “normal” age-related changes serve as the foundation that supports decreased plasticity and limited ability for tissue remodeling during pathophysiological states such as myocardial ischemia and heart failure. This review will identify populations at greater risk for poor tissue remodeling in advanced age along with present and future therapeutic strategies that may ameliorate dysfunctional tissue remodeling in aging hearts.

55. Vernemmen, I., et al. (2020). “Reference values for 2-dimensional and M-mode echocardiography in Friesian and Warmblood horses.” J Vet Intern Med 34(6): 2701–2709. BACKGROUND: Echocardiographic reference intervals for Friesian horses are poorly described. OBJECTIVES: To obtain reference intervals for echocardiographic measurements in Friesians and compare these with Warmbloods. ANIMALS: One hundred healthy adult Friesians and 100 healthy adult Warmblood horses. METHODS: Cross-sectional study. Two-dimensional and M-mode echocardiographic images were obtained. Echocardiographic measurements, including size, area, and volumetric measurements of left atrium, left and right ventricle, aorta, and pulmonary artery, were performed. Measurements were compared between the 2 breeds using an independent samples t test with Bonferroni correction for multiple comparisons. RESULTS: Reference ranges for standard echocardiographic measurements in Friesians were obtained. Several left ventricular measurements were significantly smaller in Friesians compared to Warmbloods, such as the left ventricular end-diastolic volume using the 4-chamber modified Simpsons’ method (99.85% confidence interval for the difference [CI] = –245 to –63). Also the right ventricular end-diastolic and peak-systolic internal diameter were smaller in Friesians (99.85% CI = –1.33 to –0.6 and 99.85% CI = –1.54 to –0.76, respectively). Fractional shortening (99.85% CI = 0.61-6) and ejection fraction (99.85% CI = 0.21-4.6) were significantly larger. No structural effects of systemic hypertension, such as concentric hypertrophy, were detected. CONCLUSIONS AND CLINICAL IMPORTANCE: Our study provides reference intervals for echocardiographic measurements in Friesians useful in a clinical setting. In general, the left ventricular dimensions in Friesians were significantly smaller compared to Warmbloods, emphasizing the need for breed-specific reference intervals.

56. Vinhas, M., et al. (2013). “Transthoracic echocardiography reference values in juvenile and adult 129/Sv mice.” Cardiovasc Ultrasound 11: 12. BACKGROUND: In the recent years, the use of Doppler-echocardiography has become a standard non-invasive technique in the analysis of cardiac malformations in genetically modified mice. Therefore, normal values have to be established for the most commonly used inbred strains in whose genetic background those mutations are generated. Here we provide reference values for transthoracic echocardiography measurements in juvenile (3 weeks) and adult (8 weeks) 129/Sv mice. METHODS: Echocardiographic measurements were performed using B-mode, M-mode and Doppler-mode in 15 juvenile (3 weeks) and 15 adult (8 weeks) mice, during isoflurane anesthesia. M-mode measurements variability of left ventricle (LV) was determined. RESULTS: Several echocardiographic measurements significantly differ between juvenile and adult mice. Most of these measurements are related with cardiac dimensions. All B-mode measurements were different between juveniles and adults (higher in the adults), except for fractional area change (FAC). Ejection fraction (EF) and fractional shortening (FS), calculated from M-mode parameters, do not differ between juvenile and adult mice. Stroke volume (SV) and cardiac output (CO) were significantly different between juvenile and adult mice. SV was 31.93 ± 8.67 μl in juveniles vs 70.61 ± 24.66 μl in adults, ρ < 0.001. CO was 12.06 ± 4.05 ml/min in juveniles vs 29.71 ± 10.13 ml/min in adults, ρ < 0.001. No difference was found in mitral valve (MV) and tricuspid valve (TV) related parameters between juvenile and adult mice. It was demonstrated that variability of M-mode measurements of LV is minimal. CONCLUSIONS: This study suggests that differences in cardiac dimensions, as wells as in pulmonary and aorta outflow parameters, were found between juvenile and adult mice. However, mitral and tricuspid inflow parameters seem to be similar between 3 weeks and 8 weeks mice. The reference values established in this study would contribute as a basis to future studies in post-natal cardiovascular development and diagnosing cardiovascular disorders in genetically modified mouse mutant lines.

57. Wehner, G. J., et al. (2020). “Routinely reported ejection fraction and mortality in clinical practice: where does the nadir of risk lie?” Eur Heart J 41(12): 1249–1257. AIMS: We investigated the relationship between clinically assessed left ventricular ejection fraction (LVEF) and survival in a large, heterogeneous clinical cohort. METHODS AND RESULTS: Physician-reported LVEF on 403 977 echocardiograms from 203 135 patients were linked to all-cause mortality using electronic health records (1998-2018) from US regional healthcare system. Cox proportional hazards regression was used for analyses while adjusting for many patient characteristics including age, sex, and relevant comorbidities. A dataset including 45 531 echocardiograms and 35 976 patients from New Zealand was used to provide independent validation of analyses. During follow-up of the US cohort, 46 258 (23%) patients who had undergone 108 578 (27%) echocardiograms died. Overall, adjusted hazard ratios (HR) for mortality showed a u-shaped relationship for LVEF with a nadir of risk at an LVEF of 60-65%, a HR of 1.71 [95% confidence interval (CI) 1.64-1.77] when ≥70% and a HR of 1.73 (95% CI 1.66-1.80) at LVEF of 35-40%. Similar relationships with a nadir at 60-65% were observed in the validation dataset as well as for each age group and both sexes. The results were similar after further adjustments for conditions associated with an elevated LVEF, including mitral regurgitation, increased wall thickness, and anaemia and when restricted to patients reported to have heart failure at the time of the echocardiogram. CONCLUSION: Deviation of LVEF from 60% to 65% is associated with poorer survival regardless of age, sex, or other relevant comorbidities such as heart failure. These results may herald the recognition of a new phenotype characterized by supra-normal LVEF.

58. Williams, S. L. R. E. a. A., Ed. (2012). Eurachem/CITAC guide: Quantifying Uncertainty in Analytical Measurement. www.eurachem.org.

59. Withaar, C., et al. (2021). “Heart failure with preserved ejection fraction in humans and mice: embracing clinical complexity in mouse models.” Eur Heart J 42(43): 4420–4430. Heart failure (HF) with preserved ejection fraction (HFpEF) is a multifactorial disease accounting for a large and increasing proportion of all clinical HF presentations. As a clinical syndrome, HFpEF is characterized by typical signs and symptoms of HF, a distinct cardiac phenotype and raised natriuretic peptides. Non-cardiac comorbidities frequently co-exist and contribute to the pathophysiology of HFpEF. To date, no therapy has proven to improve outcomes in HFpEF, with drug development hampered, at least partly, by lack of consensus on appropriate standards for pre-clinical HFpEF models. Recently, two clinical algorithms (HFA-PEFF and H2FPEF scores) have been developed to improve and standardize the diagnosis of HFpEF. In this review, we evaluate the translational utility of HFpEF mouse models in the context of these HFpEF scores. We systematically recorded evidence of symptoms and signs of HF or clinical HFpEF features and included several cardiac and extra-cardiac parameters as well as age and sex for each HFpEF mouse model. We found that most of the pre-clinical HFpEF models do not meet the HFpEF clinical criteria, although some multifactorial models resemble human HFpEF to a reasonable extent. We therefore conclude that to optimize the translational value of mouse models to human HFpEF, a novel approach for the development of pre-clinical HFpEF models is needed, taking into account the complex HFpEF pathophysiology in humans.

60. Xie, K., et al. (2022). “Deep phenotyping and lifetime trajectories reveal limited effects of longevity regulators on the aging process in C57BL/6J mice.” Nature Communications 13(1): 6830. Current concepts regarding the biology of aging are primarily based on studies aimed at identifying factors regulating lifespan. However, lifespan as a sole proxy measure for aging can be of limited value because it may be restricted by specific pathologies. Here, we employ large-scale phenotyping to analyze hundreds of markers in aging male C57BL/6J mice. For each phenotype, we establish lifetime profiles to determine when age-dependent change is first detectable relative to the young adult baseline. We examine key lifespan regulators (putative anti-aging interventions; PAAIs) for a possible countering of aging. Importantly, unlike most previous studies, we include in our study design young treated groups of animals, subjected to PAAIs prior to the onset of detectable age-dependent phenotypic change. Many PAAI effects influence phenotypes long before the onset of detectable age-dependent change, but, importantly, do not alter the rate of phenotypic change. Hence, these PAAIs have limited effects on aging.

61. Zacchigna, S., et al. (2021). “Towards standardization of echocardiography for the evaluation of left ventricular function in adult rodents: a position paper of the ESC Working Group on Myocardial Function.” Cardiovasc Res 117(1): 43–59. Echocardiography is a reliable and reproducible method to assess non-invasively cardiac function in clinical and experimental research. Significant progress in the development of echocardiographic equipment and transducers has led to the successful translation of this methodology from humans to rodents, allowing for the scoring of disease severity and progression, testing of new drugs, and monitoring cardiac function in genetically modified or pharmacologically treated animals. However, as yet, there is no standardization in the procedure to acquire echocardiographic measurements in small animals. This position paper focuses on the appropriate acquisition and analysis of echocardiographic parameters in adult mice and rats, and provides reference values, representative images, and videos for the accurate and reproducible quantification of left ventricular function in healthy and pathological conditions.

62. Zhang, X., et al. (2023). “How to think clearly about the central limit theorem.” Psychol Methods 28(6): 1427–1445. The central limit theorem (CLT) is one of the most important theorems in statistics, and it is often introduced to social sciences researchers in an introductory statistics course. However, the recent replication crisis in the social sciences prompts us to investigate just how common certain misconceptions of statistical concepts are. The main purposes of this article are to investigate the misconceptions of the CLT among social sciences researchers and to address these misconceptions by clarifying the definition and properties of the CLT in a manner that is approachable to social science researchers. As part of our article, we conducted a survey to examine the misconceptions of the CLT among graduate students and researchers in the social sciences. We found that the most common misconception of the CLT is that researchers think the CLT is about the convergence of sample data to the normal distribution. We also found that most researchers did not realize that the CLT applies to both sample means and sample sums, and that the CLT has implications for many common statistical concepts and techniques. Our article addresses these misconceptions of the CLT by explaining the preliminaries needed to understand the CLT, introducing the formal definition of the CLT, and elaborating on the implications of the CLT. We hope that through this article, researchers can obtain a more accurate and nuanced understanding of how the CLT operates as well as its role in a variety of statistical concepts and techniques. (PsycInfo Database Record (c) 2024 APA, all rights reserved).

