## Supplemental Data for "Establishing Comprehensive Transthoracic Echocardiography Reference Ranges for Mouse Models: Insights into the Impact of Anesthesia, Sex, and Age"

**Supplemental Table 1**: Definition and unit of measure for each TTE parameter reported.

| Parameter | Description | Unit |
|---|---|---|
| CO | Cardiac output | ml/min |
| EF | Ejection fraction | % |
| LVIDd | Left ventricular internal diameter in diastole | mm |
| LVIDs | Left ventricular internal diameter in systole | mm |
| FS | Fractional shortening | % |
| HR | Heart rate | bpm |
| LVAWd | Left ventricular anterior wall in diastole | mm |
| LVAWs | Left ventricular anterior wall in systole | mm |
| LVPWd | Left ventricular posterior wall in diastole | mm |
| LVPWs | Left ventricular posterior wall in systole | mm |
| RR | Respiration rate | l/min |
| SV | Stroke volume | µ |
| BWT | Body weight | gr |
| | Body temperature | °C |

*Supplemental Table 2*

Comprehensive overview of mean, standard deviation, and sample number for each of the nine selected TTE parameters, stratified by conscious state (conscious, anesthetized with isoflurane, or anesthetized with tribromoethanol), age [early (EA) and late (LA) adult time point] and sex (Panel a. Females; Panel b. Males).

**a**

| | Conscious - Females | | Isoflurane - Females | | Tribromoethanol - Females | |
|---|---|---|---|---|---|---|
| | EA | LA | EA | LA | EA | LA |
| *Parameter* | mean ± sd (n) | mean ± sd (n) | mean ± sd (n) | mean ± sd (n) | mean ± sd (n) | |
| *Cardiac Output [ml/min]* | 14.4 ± 3.9 (2660) | 20.6 ± 5 (378) | 16.5 ± 3.5 (3774) | 18.5 ± 4.4 (477) | no data available | no data available |
| *Ejection Fraction [%]* | 86.2 ± 7.7 (2681) | 80.1 ± 9.2 (378) | 50.4 ± 9.3 (3914) | 59.7 ± 10.3 (477) | no data available | |
| *Fractional Shortening [%]* | 55.2 ± 8.9 (2681) | 48.9 ± 9.4 (378) | 26.5 ± 6.4 (4633) | 31.4 ± 7.5 (496) | 27.9 ± 5.9 (227) | |
| *Heart Rate [bpm]* | 640.1 ± 107.3 (2681) | 593.3 ± 114.6 (378) | 452 ± 56.1 (4628) | 436.2 ± 78.9 (496) | 401.4 ± 58 (227) | |
| *LVIDd [mm]* | 2.6 ± 0.4 (2681) | 3.3 ± 0.5 (378) | 4 ± 0.3 (4631) | 4 ± 0.3 (496) | 3.9 ± 0.3 (227) | |
| *LVIDs [mm]* | 1.3 ± 0.4 (2681) | 1.7 ± 0.5 (378) | 2.9 ± 0.4 (4632) | 2.8 ± 0.5 (496) | 2.8 ± 0.4 (227) | |
| *LVPWd [mm]* | 0.5 ± 0 (2681) | 0.5 ± 0 (378) | 0.7 ± 0.1 (4632) | 0.8 ± 0.1 (496) | 0.6 ± 0.1 (227) | |
| *LVPWs [mm]* | 0.5 ± 0 (2681) | 0.5 ± 0 (378) | 0.9 ± 0.2 (3913) | 1.1 ± 0.2 (477) | no data available | |
| *Stroke Volume [µl]* | 23.1 ± 7.1 (2667) | 36.1 ± 10.9 (378) | 35.6 ± 6.2 (3789) | 42.5 ± 7.4 (477) | no data available | |

**b**

| | Conscious - Males | | Isoflurane - Males | | Tribromoethanol - Males | |
|---|---|---|---|---|---|---|
| | EA | LA | EA | LA | EA | LA |
| *Parameter* | mean ± sd (n) | mean ± sd (n) | mean ± sd (n) | mean ± sd (n) | mean ± sd (n) | |
| *Cardiac Output [ml/min]* | 16.3 ± 4.2 (2675) | 21.8 ± 6 (376) | 18.9 ± 4.1 (2695) | 20.2 ± 4.7 (413) | no data available | no data available |
| *Ejection Fraction [%]* | 84.8 ± 7.6 (2694) | 82.5 ± 8.2 (376) | 49.5 ± 9.4 (2841) | 51.4 ± 9.1 (413) | no data available | |
| *Fractional Shortening [%]* | 53.5 ± 8.3 (2694) | 51.3 ± 8.9 (376) | 26 ± 6.3 (3590) | 26.1 ± 6 (432) | 27.1 ± 7.2 (223) | |
| *Heart Rate [bpm]* | 654.2 ± 98.9 (2694) | 648.1 ± 94.8 (376) | 457.7 ± 64.2 (3588) | 415.7 ± 67.9 (432) | 394.2 ± 59.9 (223) | |
| *LVIDd [mm]* | 2.8 ± 0.4 (2694) | 3.2 ± 0.5 (376) | 4.2 ± 0.4 (3588) | 4.6 ± 0.4 (432) | 4.1 ± 0.3 (223) | |
| *LVIDs [mm]* | 1.4 ± 0.4 (2694) | 1.6 ± 0.5 (376) | 3.1 ± 0.5 (3591) | 3.4 ± 0.5 (432) | 3 ± 0.4 (223) | |
| *LVPWd [mm]* | 0.5 ± 0 (2694) | 0.5 ± 0 (376) | 0.7 ± 0.1 (3590) | 0.8 ± 0.1 (432) | 0.6 ± 0.1 (223) | |
| *LVPWs [mm]* | 0.5 ± 0 (2694) | 0.5 ± 0 (376) | 1 ± 0.2 (2842) | 1.1 ± 0.2 (413) | no data available | |
| *Stroke Volume [µl]* | 25.5 ± 7.6 (2680) | 34.5 ± 11 (376) | 39.7 ± 7.2 (2703) | 48.8 ± 8.9 (413) | no data available | |

***Supplemental Table 3***

Histograms presenting the distribution of LVAWd and LWAVs data along with calculated ranges (mean ± SD and median and 95% reference range) for isoflurane anesthetized EA and LA mice, stratified by sex. These calculations are based on data from three (out of six) contributing centers.

Panel A and B depict early (EA) and Panel C and D late (LA) adult time point.

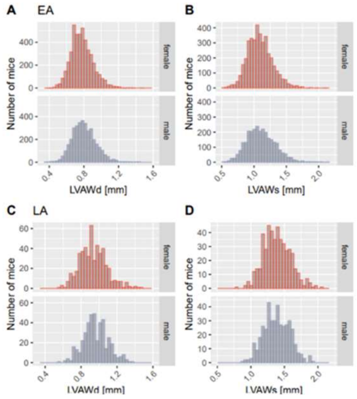

**Females**

| Parameter | EA median [95% range] | LA median [95% range] |
|---|---|---|
| *LVAWd [mm]* | 0.8 [0.6;1.1] | 0.9 [0.7;1.3] |
| *LVAWs [mm]* | 1.1 [0.7;69.6] | 1.4 [1;70.8] |

**Males**

| Parameter | EA median [95% range] | LA median [95% range] |
|---|---|---|
| *LVAWd [mm]* | 0.8 [0.6;1.1] | 0.9 [0.7;1.3] |
| *LVAWs [mm]* | 1.1 [0.7;69.6] | 1.4 [1;70.8] |

**Females**

| Parameter | EA mean ± sd (n) | LA mean ± sd (n) |
|---|---|---|
| *LVAWd [mm]* | 0.8 ± 0.1 (4633) | 0.9 ± 0.1 (496) |
| *LVAWs [mm]* | 1.1 ± 0.2 (3914) | 1.4 ± 0.2 (477) |

**Males**

| Parameter | EA mean ± sd (n) | LA mean ± sd (n) |
|---|---|---|
| *LVAWd [mm]* | 0.8 ± 0.1 (3590) | 1 ± 0.1 (432) |
| *LVAWs [mm]* | 1.1 ± 0.2 (2842) | 1.4 ± 0.2 (413) |

***Supplemental Figure 1***

Histograms presenting the distribution of body temperature data along with calculated ranges (mean ± SD and median and 95% reference range) for isoflurane anesthetized EA and LA mice, stratified by sex. These calculations are based on data from five (out of six) contributing centers.

Panel A and B depict early (EA) and Panel C and D late (LA) adult time point.

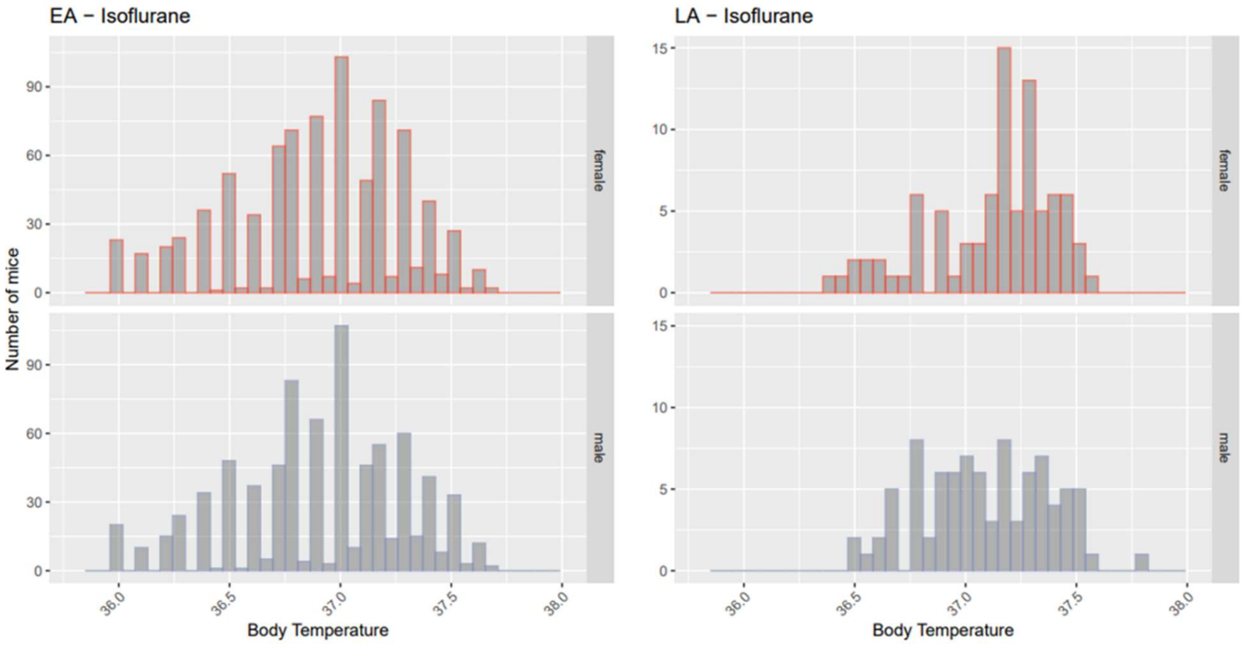

| | mean ± sd (n) | median [95% range] |
|---|---|---|
| *Body Temperature – female* | 36.9 ± 0.38 (854) | 36.9 [36;37.5] |
| *Body Temperature – male* | 36.92 ± 0.38 (803) | 37 [36.1;37.5] |
| *Body Temperature – both* | 36.91 ± 0.38 (1657) | 36.9 [36;37.5] |

| mean ± sd (n) | median [95% range] |
|---|---|
| 37.13 ± 0.28 (88) | 37.2 [36.5;37.5] |
| 37.09 ± 0.29 (88) | 37.1 [36.6;37.5] |
| 37.11 ± 0.28 (176) | 37.2 [36.5;37.5] |

***Supplemental Figure 2***

Testing sex-differences in conscious mice.
T-test results when comparing data from conscious male and female animals for each of the nine selected TTE parameters, stratified by age [early (EA) and late (LA) adult time point]) demonstrated that some parameters show high significance (p<.001), while for others there was no evidence of sexual dimorphism (LVPWd, LVPWs in EA and SV in LA). Associated Cohen´s d standardized effect sizes were negligible to small and for LA HR medium.

Panel a shows EA time point, Panel b shows LA time point.

**a** Early adult (EA)

**Conscious – SEX EFFECT**

| | p value | Cohen's D |
|---|---|---|
| *Cardiac Output [ml/min]* | p<.001 | -0.48 (small) |
| *Ejection Fraction [%]* | p<.001 | 0.17 (negligible) |
| *Fractional Shortening [%]* | p<.001 | 0.2 (negligible) |
| *Heart Rate [bpm]* | p<.001 | -0.14 (negligible) |
| *LVIDd [mm]* | p<.001 | -0.28 (small) |
| *LVIDs [mm]* | p<.001 | -0.27 (small) |
| *LVPWd [mm]* | p=.179 | -0.04 (negligible) |
| *LVPWs [mm]* | p=.015 | -0.07 (negligible) |
| *Stroke Volume [µl]* | p<.001 | -0.33 (small) |

**b** Late adult (LA)

**Conscious – SEX EFFECT**

| | p value | Cohen's D |
|---|---|---|
| *Cardiac Output [ml/min]* | p=.002 | -0.22 (small) |
| *Ejection Fraction [%]* | p<.001 | -0.27 (small) |
| *Fractional Shortening [%]* | p<.001 | -0.26 (small) |
| *Heart Rate [bpm]* | p<.001 | -0.52 (medium) |
| *LVIDd [mm]* | p=.004 | 0.21 (small) |
| *LVIDs [mm]* | p<.001 | 0.27 (small) |
| *LVPWd [mm]* | p=.1 | -0.12 (negligible) |
| *LVPWs [mm]* | p=.753 | -0.02 (negligible) |
| *Stroke Volume [µl]* | p=.052 | 0.14 (negligible) |

***Supplemental Figure 3***

Testing sex-differences in isoflurane anesthetized mice.
T-test results when comparing data from conscious male and female animals for each of the nine selected TTE parameters, stratified by age [early (EA) and late (LA) adult time point]) demonstrated that some parameters show high significance ($p<.001$), while for others there was no evidence of sexual dimorphism (LVPWs in LA). Associated Cohen´s d standardized effect sizes were negligible to small and for CO, LVIDd and SV medium in EA. EF, LVIDd, LVIDs showed large and FS and SV medium effect sizes in LA.

Panel a shows EA time point, Panel b shows LA time point.

**a** Early adult (EA)

**Isoflurane - SEX**

| | p value | Cohen's D |
|---|---|---|
| *Cardiac Output [ml/min]* | p<.001 | -0.63 (medium) |
| *Ejection Fraction [%]* | p<.001 | 0.1 (negligible) |
| *Fractional Shortening [%]* | p<.001 | 0.08 (negligible) |
| *Heart Rate [bpm]* | p<.001 | -0.1 (negligible) |
| *LVIDd [mm]* | p<.001 | -0.56 (medium) |
| *LVIDs [mm]* | p<.001 | -0.38 (small) |
| *LVPWd [mm]* | p<.001 | -0.34 (small) |
| *LVPWs [mm]* | p<.001 | -0.35 (small) |
| *Stroke Volume [µl]* | p<.001 | -0.62 (medium) |

**b** Late adult (LA)

**Isoflurane - SEX**

| | p value | Cohen's D |
|---|---|---|
| *Cardiac Output [ml/min]* | p<.001 | -0.37 (small) |
| *Ejection Fraction [%]* | p<.001 | 0.85 (large) |
| *Fractional Shortening [%]* | p<.001 | 0.77 (medium) |
| *Heart Rate [bpm]* | p<.001 | 0.28 (small) |
| *LVIDd [mm]* | p<.001 | -1.51 (large) |
| *LVIDs [mm]* | p<.001 | -1.27 (large) |
| *LVPWd [mm]* | p<.001 | -0.35 (small) |
| *LVPWs [mm]* | p=.926 | -0.01 (negligible) |
| *Stroke Volume [µl]* | p<.001 | -0.78 (medium) |

***Supplemental Figure 4***

Testing sex-differences in tribromoethanol anesthetized mice.

T-test results when comparing data from conscious male and female animals for each of the nine selected TTE parameters, stratified by early (EA) adult time point demonstrated that some parameters show high significance (p<.001), while for others there was no evidence of sexual dimorphism (HR and FS). Associated Cohen´s d standardized effect sizes were negligible to small and for LVIDd in EA.

Panel a shows EA time point.

**a** Early adult (EA)

**Tribromoethanol - SEX**

| | p value | Cohen's D |
|---|---|---|
| *Fractional Shortening [%]* | p=.171 | 0.13 (negligible) |
| *Heart Rate [bpm]* | p=.2 | 0.12 (negligible) |
| *LVIDd [mm]* | p<.001 | -0.52 (medium) |
| *LVIDs [mm]* | p<.001 | -0.4 (small) |
| *LVPWd [mm]* | p=.006 | -0.26 (small) |

***Supplemental Figure 5:*** Seven inbred strains of The Jaxwest1 project 129S1/SvImJ, A/J, BALB/cJ, C57BL/6J, DBA/2J, NOD/ShiLtJ and SJL/J show a close alignment to the reference ranges reported herein for CO, EF, LVIDd, LVIDs, FS, and HR based on multiple C57BL/6N substrains indicating good utility for those reference ranges. Data on LVPWd, LVPWs and SV were not available in this study. Mice were isoflurane anesthetized, split by sex and ~12 weeks of age, equivalent to the IMPC EA time point.

Red dotted lines depict the boundaries of the sex-specific reference range calculated herein, for each parameter.

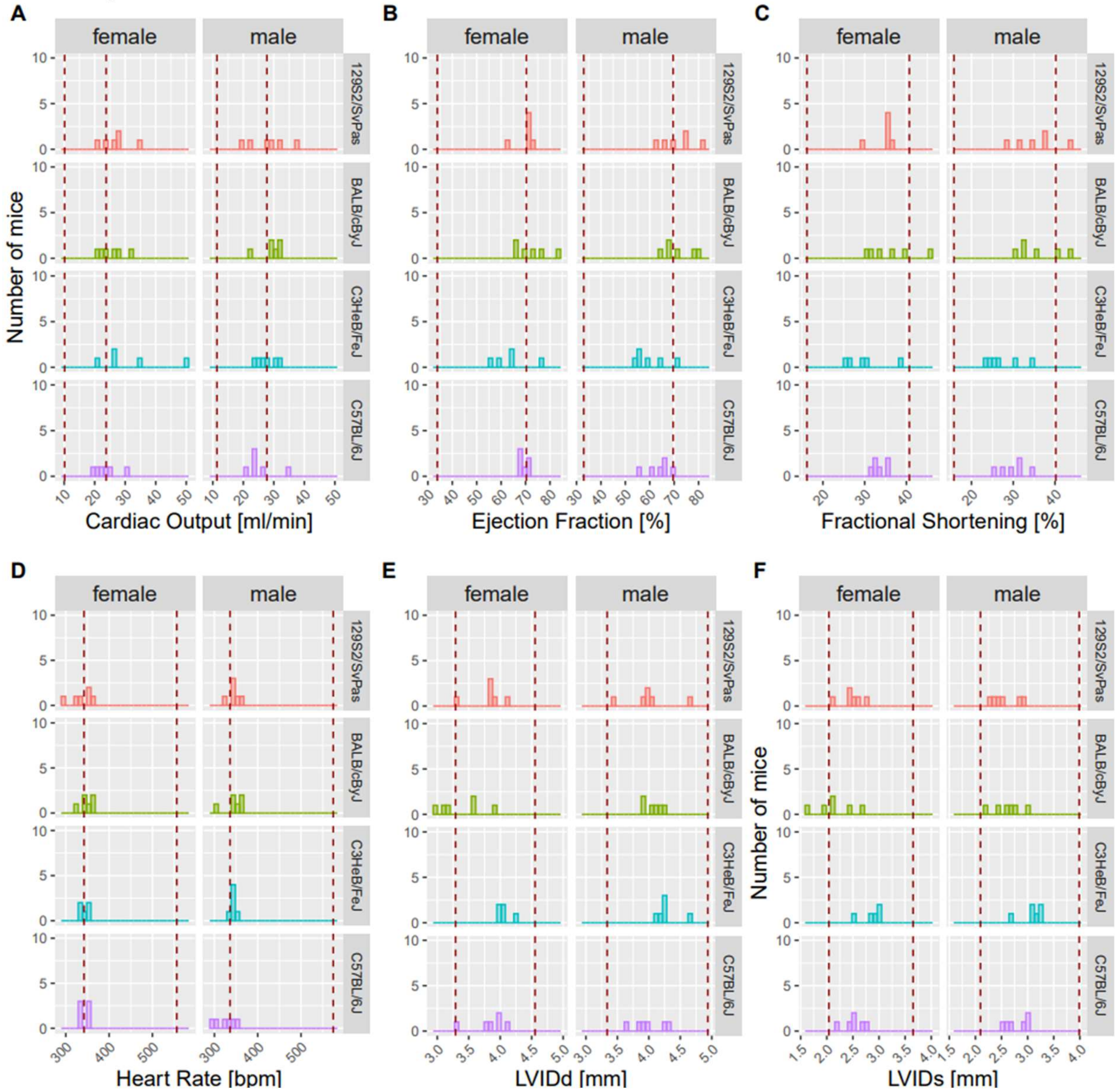

***Supplemental Figure 5:*** Seven inbred strains of The Jaxwest1 project 129S1/SvImJ, A/J, BALB/cJ, C57BL/6J, DBA/2J, NOD/ShiLtJ and SJL/J show a close alignment to the reference ranges reported herein for CO, EF, LVIDd, LVIDs, FS, HR, LVPWd, LVPWs and SV based on multiple C57BL/6N substrains indicating good utility for those reference ranges. Mice were isoflurane anesthetized, split by sex and ~12 weeks of age, equivalent to the IMPC EA time point.

Red dotted lines depict the boundaries of the sex-specific reference range calculated herein, for each parameter.

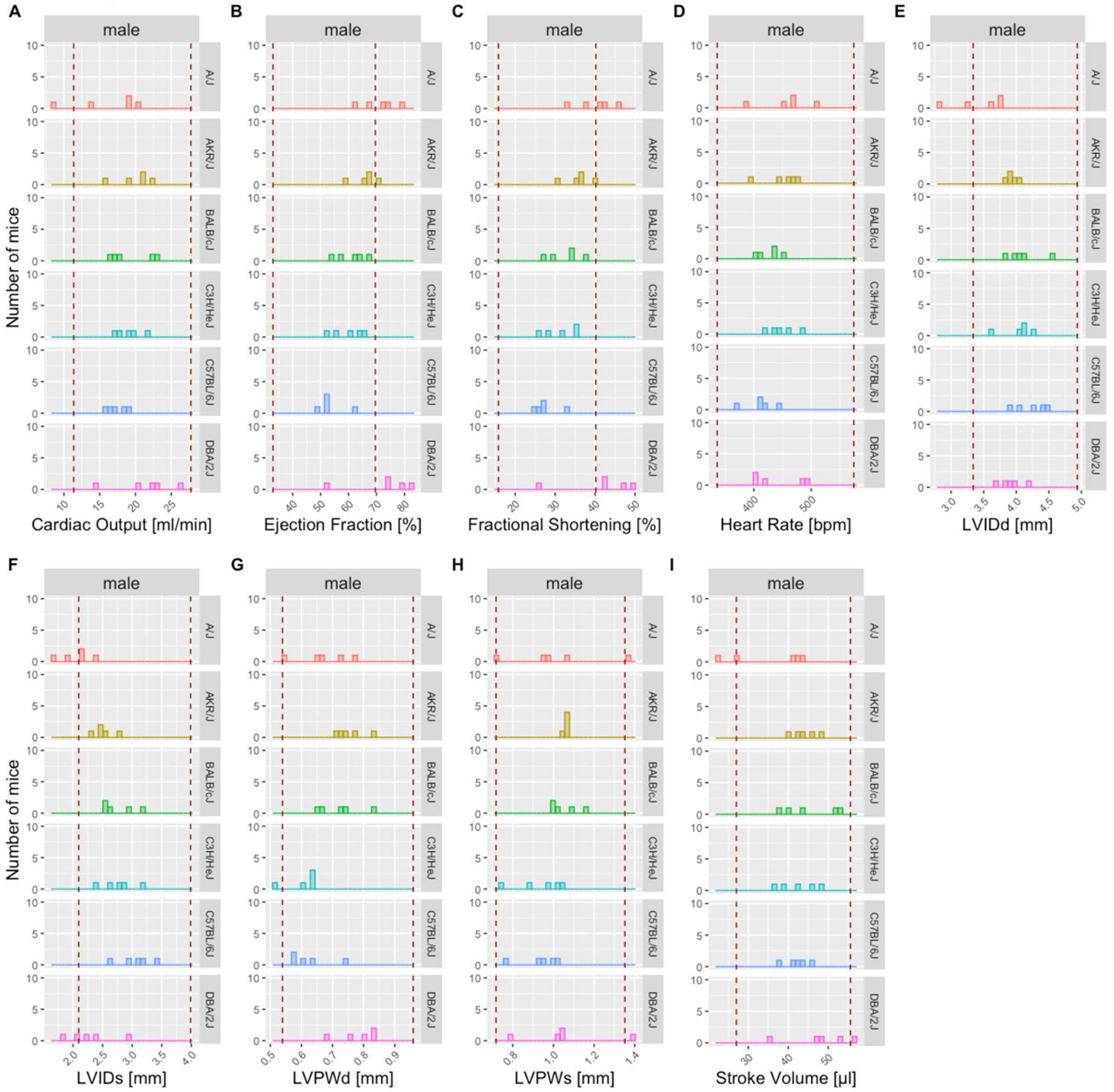

***Supplemental Figure 6:*** Three inbred, wildtype control animals of non-IMPC studies conducted at the German Mouse Clinic C57BL/6J, C57BL/6N, and FVB show a close alignment to the reference ranges reported herein for CO, EF, LVIDd, LVIDs, FS, HR, LVPWd, LVPWs and SV based on multiple C57BL/6N substrains indicating good utility for those reference ranges. Mice were conscious, split by sex and ~12 weeks of age, equivalent to the IMPC EA time point.

Red dotted lines depict the boundaries of the sex-specific reference range calculated herein, for each parameter.

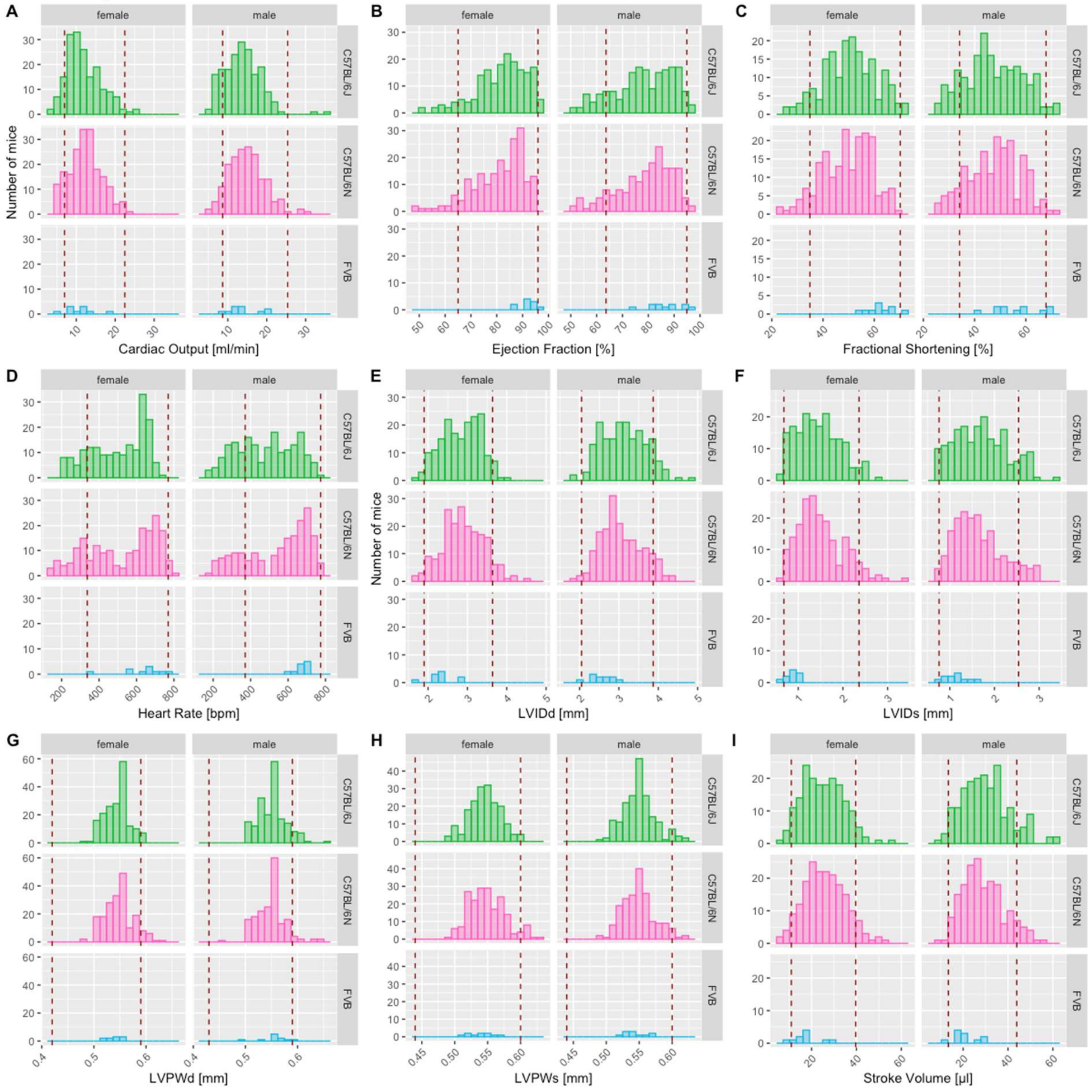
